# RTK-dependent inducible degradation of mutant PI3Kα drives GDC-0077 (Inavolisib) efficacy

**DOI:** 10.1101/2021.05.12.442687

**Authors:** Kyung W. Song, Kyle A. Edgar, Emily J. Hanan, Marc Hafner, Jason Oeh, Mark Merchant, Deepak Sampath, Michelle A. Nannini, Rebecca Hong, Lilian Phu, William F. Forrest, Eric Stawiski, Stephen Schmidt, Nicholas Endres, Jane Guan, Jeffrey J. Wallin, Jonathan Cheong, Emile Plise, Gail Philips, Laurent Salphati, Timothy P. Heffron, Alan Olivero, Shiva Malek, Steven T. Staben, Donald S. Kirkpatrick, Anwesha Dey, Lori S. Friedman

**Affiliations:** Departments of Discovery Oncology, Genentech, Inc., 1 DNA Way, South San Francisco, California 94080, USA; Departments of Discovery Chemistry, Genentech, Inc., 1 DNA Way, South San Francisco, California 94080, USA; Departments of Microchemistry, Proteomics & Lipidomics, Genentech, Inc., 1 DNA Way, South San Francisco, California 94080, USA; Departments of Bioinformatics, Genentech, Inc., 1 DNA Way, South San Francisco, California 94080, USA; Departments of Structural Biology, Genentech, Inc., 1 DNA Way, South San Francisco, California 94080, USA; Departments of Drug Metabolism and Pharmacokinetics, Genentech, Inc., 1 DNA Way, South San Francisco, California 94080, USA; Departments of Translational Oncology, Genentech, Inc., 1 DNA Way, South San Francisco, California 94080, USA; Departments of Biochemical and Cell Biology, Genentech, Inc., 1 DNA Way, South San Francisco, California 94080, USA

**Keywords:** GDC-0077 (Inavolisib), Taselisib, breast cancer, HER2-positive, PIK3CA, PI3K, p110α

## Abstract

*PIK3CA* is one of the most frequently mutated oncogenes; the p110α protein it encodes plays a central role in tumor cell proliferation and survival. Small molecule inhibitors targeting the PI3K p110α catalytic subunit have entered clinical trials, with early-phase GDC-0077 (Inavolisib) studies showing anti-tumor activity and a manageable safety profile in patients with *PIK3CA*-mutant, hormone receptor-positive breast cancer as a single agent or in combination therapy. Despite this, preclinical studies have shown that PI3K pathway inhibition releases negative feedback and activates receptor tyrosine kinase signaling, reengaging the pathway and attenuating drug activity. Here we discover that GDC-0077 and taselisib more potently inhibit mutant PI3K pathway signaling and cell viability through unique HER2-dependent degradation. Both are more effective than other PI3K inhibitors at maintaining prolonged pathway suppression, resulting in enhanced apoptosis and greater efficacy. This unique mechanism against mutant p110α reveals a new strategy for creating inhibitors that specifically target mutant tumors with selective degradation of the mutant oncoprotein and also provide a strong rationale for pursuing PI3Kα degraders in patients with HER2-positive breast cancer.

## Introduction

Oncogenic mutations in the *PIK3CA* gene increase lipid kinase activity and transform cells (Isakoff et al., 2005; Kang et al., 2005; Samuels et al., 2005). The alpha isoform of PI3K is a dimer composed of the p110α catalytic subunit and a p85 regulatory subunit which functions to stabilize p110α and reduce kinase activity (Yu et al., 1998). The binding of a phosphorylated receptor tyrosine kinase (RTK) activates p110α through the release of a subset of inhibitory contacts with p85 (Burke and Williams, 2013). Common hotspot mutations in *PIK3CA* helical (*E542K*, *E545K*) and kinase (*H1047R*) domains function by perturbing local interfaces between p85 and p110α (Echeverria et al., 2015; Miled et al., 2007) and increasing dynamic events required for catalysis on membranes (Burke et al., 2012). Several inhibitors of PI3K have entered clinical trials, yet in patients with *PIK3CA*-mutant tumors the efficacy has been modest, in part due to a limited therapeutic index (Krop et al., 2016; Martin et al., 2017; Mayer et al., 2017; Rodon et al., 2013). Hence, we reasoned that it might be possible to improve the therapeutic index by identifying compounds with increased specificity for mutant p110α.

## Results

### PI3K inhibitor potency in *PIK3CA*-mutant cells

A selection of PI3K inhibitors were profiled for biochemical activity and pharmacokinetic properties, including inhibitors across several chemical classes and with varying p110 isoform selectivity (Figure 1A). Taselisib and GDC-0077 showed increased mutant potency in cell viability assays in a cancer cell line panel compared with other PI3K inhibitors (including the alpha isoform-selective inhibitor, BYL719) (Figure 1B). Furthermore, taselisib (and later, GDC-0077) was a stronger inducer of cell death compared with other compounds, specifically in *PIK3CA*-mutant cancer cell lines (Figure S1A), suggesting that taselisib is more potent in *PIK3CA*-mutant cells compared with to other PI3K inhibitors. We compared PI3K inhibitor potencies in parental isogenic SW48 colon cancer cells bearing wild-type (WT) *PIK3CA* and matched isogenic lines expressing *H1047R* or *E545K* hotspot mutants knocked into one allele of the *PIK3CA* locus. Taselisib potency (half maximal effective concentration [EC_50_]) increased 3-fold in *PIK3CA*-mutant cells versus parental WT SW48 cells, while GDC-0941 (Folkes et al., 2008) displayed comparable EC_50_ in mutant and WT cells (Figure 1C). To assess whether this potency shift was correlated with the taselisib chemical scaffold, structurally related analogs with increased alpha-isoform specificity, GNE-102 and GNE-326 (Heffron et al., 2016), and a PI3Kα inhibitor from an unrelated chemical class (BYL719) (Furet et al., 2013) were assessed and found to not have a potency differential in mutant versus WT isogenic cells (Figure S1B). We also confirmed that inhibition of multiple PI3K isoforms did not play a role in this increased potency. Taselisib binds equipotently to p110α and p110δ isoforms but is selective against p110β and p110γ isoforms (Figure 1A). However, combination of a p110α inhibitor (GNE-102) with a p110δ inhibitor (idelalisib) (Sadhu et al., 2003) did not impact cell potency, nor did the combination of taselisib with a p110β inhibitor (TGX-221) (Jackson et al., 2005) (Figures S1C and S1D). Neither cell permeability differences nor intracellular accumulation of inhibitors could explain the increased potency of taselisib in mutant cells (Figure S1E).

**Figure 1.**
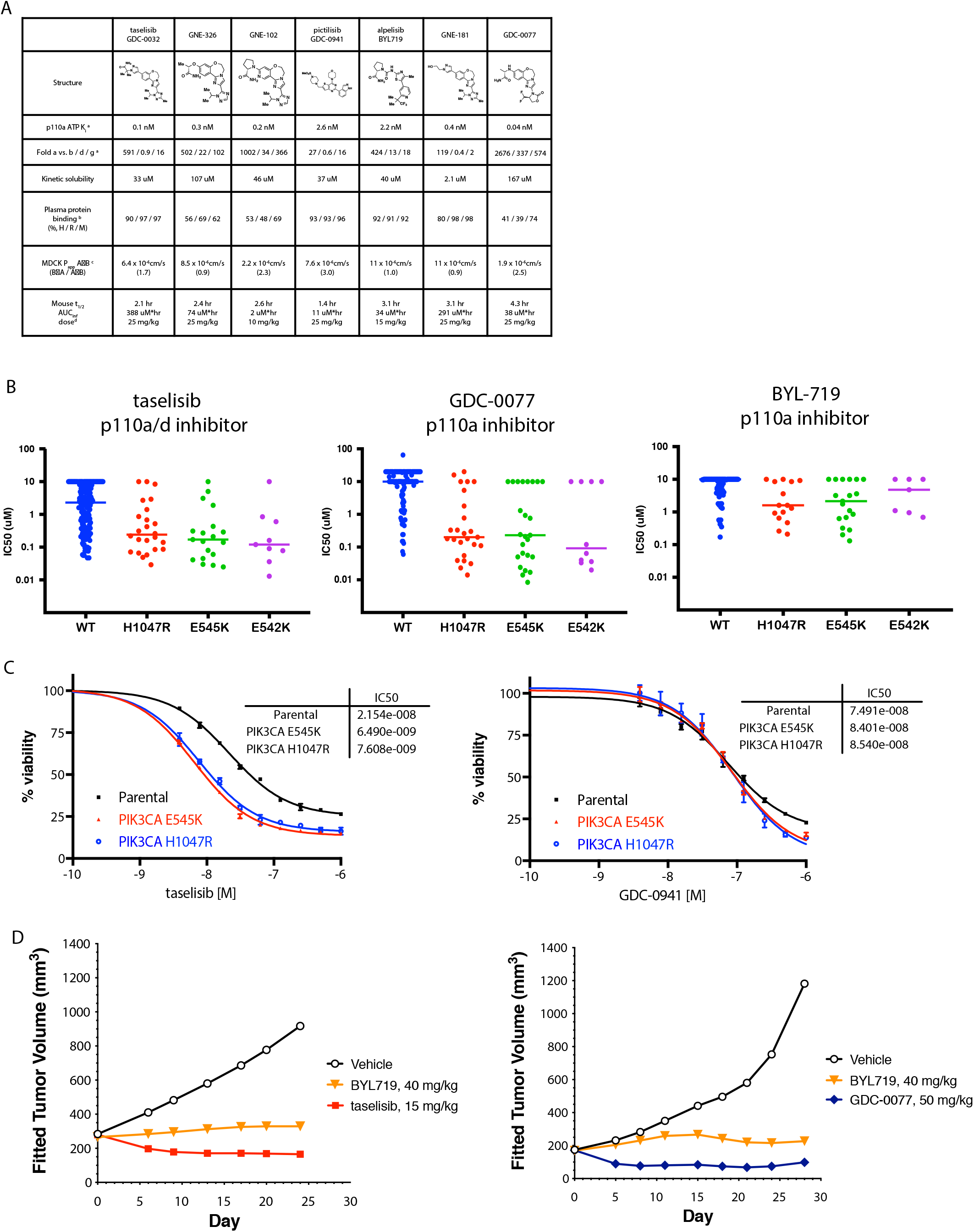
GDC-0032 and GDC-0077 have increased potency in *PIK3CA*-mutant cancer cells. **(A)** Chemical structures and physicochemical properties of PI3K inhibitors. ^a^Inhibition of ATP-hydrolysis by PI3K isoforms in a biochemical assay, ADP production measured by ADP-Glo^TM^. ^b^Plasma protein binding determined by equilibrium dialysis. ^c^ Permeability measured using Madin-Darby canine kidney (MDCK) epithelial cells; A = apical, B = basolateral. B➔A / A➔B used to estimate efflux potential.^d^ Mouse PO dose as MCT suspension. **(B)** Cell viability IC_50_ values determined by quantifying ATP from all tumor lines at 5 days post-treatment. **(C)** PI3K inhibitor potency in SW48 isogenic *PIK3CA*-mutant and -wild-type parental cells in a 4-day viability assay. Error bars are standard deviation of quadruplicates **(D)** *In vivo* efficacy of taselisib, GDC-0077, and BYL719 in HCC1954 PIK3CA H1047R breast cancer xenograft model.

### *In vivo* efficacy of GDC-0077

We next asked whether this greater potency and enhanced cell death manifested *in vivo* in mutant PI3K tumor xenografts. Indeed, we observed greater efficacy for taselisib and GDC-0077 compared with a maximum tolerated dose (MTD) of BYL719 (Figure 1D). In addition, GDC-0077 treatment at the MTD *in vivo* resulted in tumor regressions in multiple *PIK3CA*-mutant xenograft and patient-derived xenograft models (HCC1954, KPL4, and HCI-003 PDX) (Figures S1F).

Given the potency of GDC-0077 and the significant improvement in PI3Kα isoform selectivity over both taselisib and BYL719 (Figures 1A and 1B), we next evaluated the efficacy of GDC-0077 and the suitability for combination with standard of care in hormone receptor (HR)-positive/HER2-negative breast cancers. These included aromatase inhibitors and, more recently, CDK4/6 inhibitors such as palbociclib. We therefore assessed the effect of combining GDC-0077 with these drugs to evaluate efficacy and safety. First, we measured *in vitro* growth of five *PI3KCA*-mutant HR-positive lines across different concentrations of GDC-0077 with or without E2 (to mimic aromatase inhibitors [AIs]) and with or without 0.15 *µ*M of palbociclib. To assess growth, endpoint cell population, as measured by CyQuant assay, was normalized to the cell population at the time of treatment using the growth-rate inhibition (GR) method (Hafner et al., 2016). A GR value of 1 meant no inhibition, 0 meant no net growth, and negative values represented cell loss. Response in *PIK3CA E545K*-mutant MCF-7 cells showed that addition of GDC-0077 induced a strong cytotoxic response in all combination treatments, as reflected by negative GR values for concentrations of 0.12*µ*M and above (Hafner et al., 2016) (Figure 2A). When assessing the efficacy of the combination treatments across all five cell lines, we found that the GR values measured in the condition with palbociclib and without E2 (equivalent to AI) decreased by a median of 0.25 when adding 0.123 *µ*M of GDC-0077, showing a broad increase in efficacy by combining GDC-0077 with standard of care treatments (Figure 2B and S2A).

**Figure 2.**
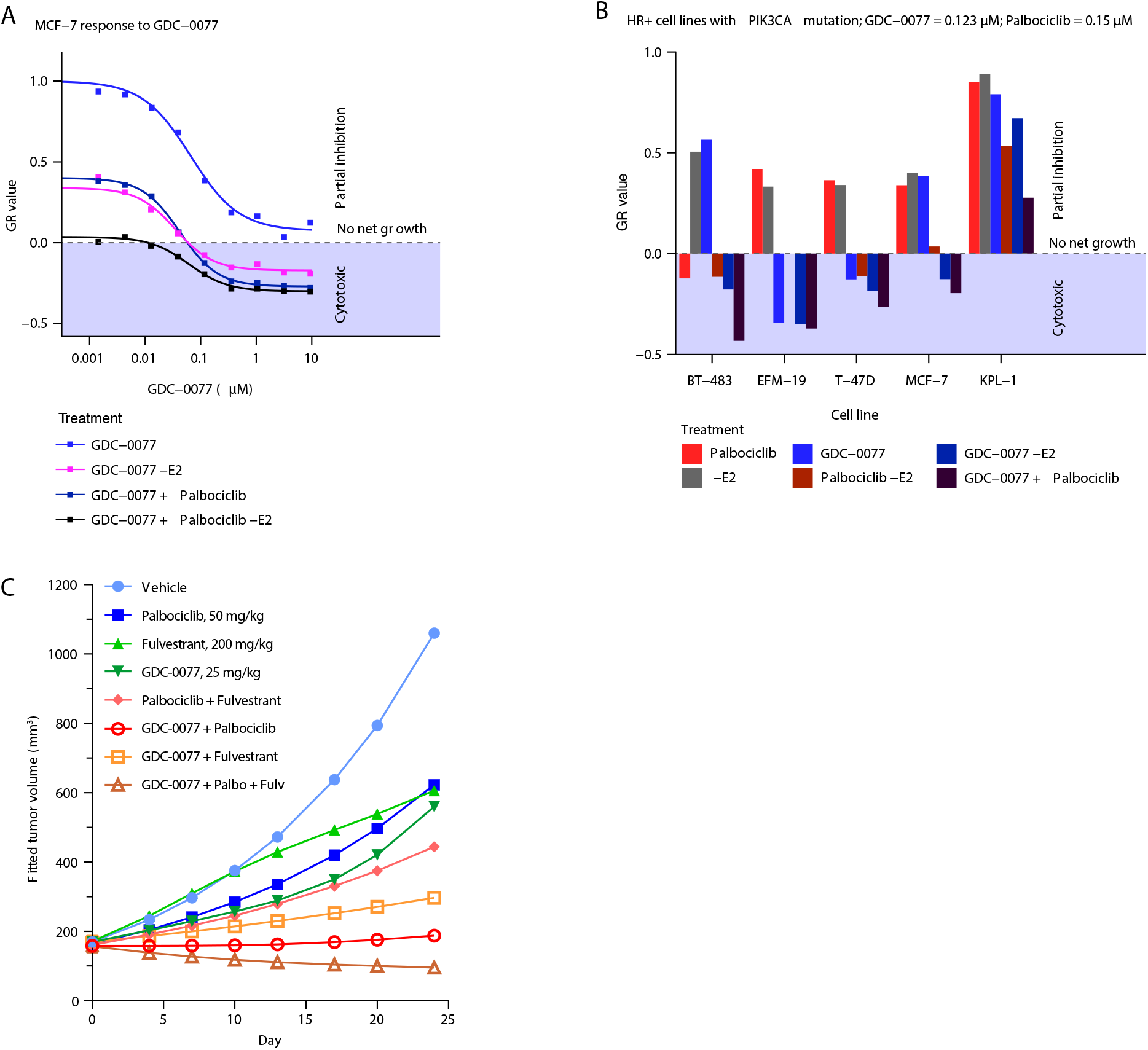
Activity of PI3Kα inhibitors in combination with palbociclib in HR-positive breast cancer cells. **(A)** Dose response curve of MCF-7 cells treated with GDC-0077 either alone (blue), without E2 to mimic aromatase inhibitor (dark blue), with palbociclib at 0.15*µ*M (purple), or with palbociclib at 0.15 *µ*M (purple) and without E2 (dark purple). Negative values indicate a cytotoxic response. y-axis shows normalized growth inhibition (GR value). **(B)** Growth response of 5 HR-positive breast cancer cells with *PIK3CA* mutations to treatment with either palbociclib at 0.15 *µ*M (red), E2 withdrawal (gray), GDC-0077 at 0.123 *µ*M (blue), Palbociclib at 0.15*µ*M and E2 withdrawal (dark red), or the triple combination (dark purple). y-axis shows normalized growth inhibition (GR value). **(C)** Efficacy of palbociclib, fulvestrant and GDC-0077 as single agent or in combination in MCF7 xenografts. Each group contains 12 animals at the beginning of the experiment.

We then sought to validate this increased efficacy *in vivo*. Palbociclib (50 mg/kg) combined with fulvestrant (200mg/kg) only conferred tumor growth inhibition (TGI) of 71%, whereas addition of GDC-0077 (25 mg/kg) further reduced tumor burden (mean TGI of 106%, N = 12, Figure 2C), consistent with the response observed *in vitro* (Figure 2A). Weight loss was modest at 8.1% in the triple combination cohort (Figure S2B), suggesting tolerability of adding GDC-0077 to a regimen with endocrine therapy plus CDK4/6 inhibitors. Similar efficacy results were obtained with taselisib combined with palbociclib (Figure S2C), although weight loss was higher (Figure S2C) with taselisib as expected. Also, taselisib could only be combined with palbociclib and not the triple combination. Taken together, these data suggested the possibility of combining GDC-0077 with both palbociclib and fulvestrant for superior efficacy in *PIK3CA*-mutant, *HER2*-negative tumors and providing a therapeutic window not achievable with earlier PI3K inhibitors.

### Taselisib and GDC-0077 induce mutant p110α degradation

In order to determine the mechanistic basis for this efficacy differential in *PIK3CA*-mutant tumors, we compared the effects downstream (pAKT) and upstream (pHER3) of PI3K signaling for GDC-0077 versus BYL719 over a timecourse. In both cases we observed robust acute inhibition of pAKT treatment; however, GDC-0077 demonstrated sustained inhibition of pAKT over the 24-hour treatment time despite inducing release of negative feedback, as measured by upregulation of pHER3 (as is also observed for BYL719) (Figure 3A).

**Figure 3.**
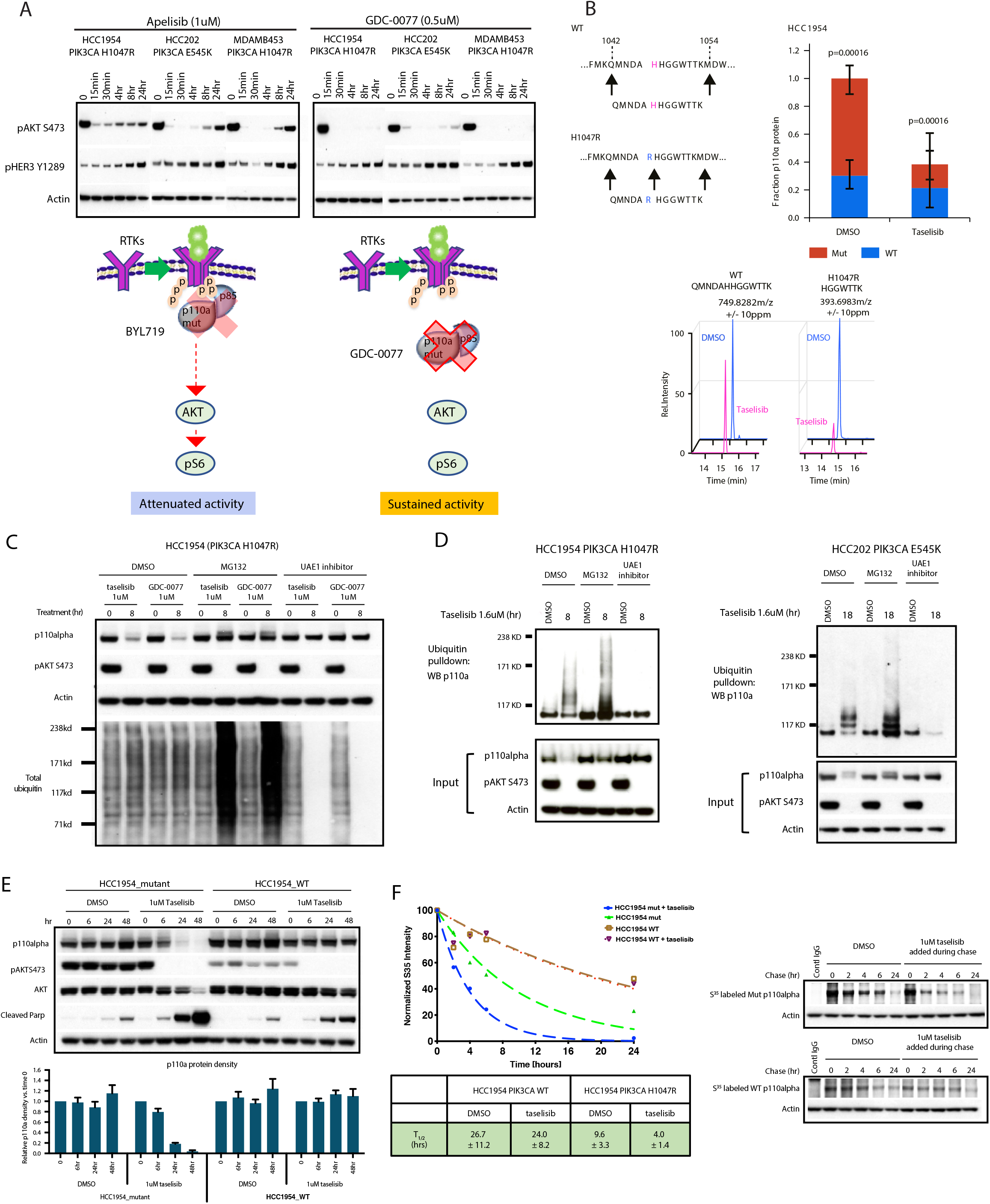
Taselisib depletes mutant p110α protein through ubiquitin and proteasome mechanism in a dose and time dependent manner. **(A)** Western blots of the inhibitor response in PI3K signaling (pHER3 and pAKT) in *PIK3CA*-mutant (HCC1954, HCC202, and MDA-MB-453) cell lines treated with 1 *µ*M BYL719 or 0.5 *µ*M GDC-0077 for indicated time points. **(B)** Mass spectrometry of HCC1954 cells treated for 24 hours with 500 nM taselisib. A neo-tryptic peptide generated from PIK3CA H1047R was used to assess mutant protein levels compared to wild-type protein in the same lysate. **(C)** Rescue of taselisib- or GDC-0077-mediated p110α degradation in HCC1954 cells by either a proteasome inhibitor (MG132) or a Ubiquitin Activating Enzyme E1 (UAE1) inhibitor. **(D)** HCC1954 PIK3CA H1047R cells were treated 8 hours or HCC202 PIK3CA E545K cells were treated for 18 hours with either DMSO or 1 *µ*M taselisib ± MG132 or ± UAE1 inhibitor. Protein lysates were run on western blot and probed with antibodies to p110α, pAKT, and b-actin, or ubiquitinated proteins were pulled-down with TUBE1 reagent and then blotted with anti-p110α antibody. **(E)** HCC1954 cells engineered to be isogenic for *PIK3CA*-mutant or -wild-type were treated with 1 μM taselisib for up to 48 hrs followed by western blotting. Experimental replicates n=3 were used to quantify p110α. **(F)** Pulse-chase of isogenic cell lines, HCC1954_mutant and HCC1954_wild-type. PI3K inhibitor taselisib at 1 *µ*M was added during the chase. Pull-down with p110α antibody was followed by autoradiography, and data fit to a single exponential decay function.

We reasoned that one possible mechanism of enabling of sustained inhibition of pAKT in spite of elevated pHER3 levels would be via drug-induced sequestration of PI3K away from the plasma membrane. In order to investigate this hypothesis, levels of p110α protein in sub-cellular fractions were evaluated by western blot at various timepoints after taselisib treatment. Unexpectedly, we discovered time-dependent p110α protein depletion from the *PIK3CA*-mutant cells following taselisib treatment, regardless of the subcellular fraction evaluated (Figure S3A). BYL719 did not significantly impact p110α protein levels. Comparing whole cell lysates from *PIK3CA*-mutant and -WT breast cancer cells by western blot, taselisib treatment for 8 hours caused the dose-dependent depletion of p110α protein specifically in *PIK3CA H1047R*-mutant HCC1954 cells. No significant change was observed for p110α in *PIK3CA*-WT HDQ-P1 cells (Figure S3B). Treatment with BYL719 did not affect p110α levels in any cell lines tested. Transcription of *PIK3CA* alleles was not diminished following taselisib treatment (Figure S3C), implicating post-transcriptional regulation of mutant p110α protein.

We investigated p110α protein depletion in p110α mutant xenograft breast cancer models and found that membrane-associated p110α was depleted 4 hours after a single oral dose of 15 mg/kg taselisib in individual tumors from a p110α-mutant HCC1954 xenograft model, and that p110α depletion was not observed with 40 mg/kg BYL719 (Figure S3D). In the same xenograft model, a single oral dose of 50 mg/kg GDC-0077 depleted p110α protein expression for up to 8 hours, further confirming a similar mechanism of action for both taselisib and GDC-0077. In Phase Ia clinical trials, taselisib and GDC-0077 had antitumor activity in *PIK3CA*-mutant tumors as assessed by response rates (Juric et al., 2017). The free drug exposures achieved in the clinic were evaluated in tissue culture experiments with *PIK3CA* mutant breast cancer lines. Unlike pictilisib (GDC-0941), both taselisib (GDC-0032) and inavolisib (GDC-0077) treatment for 24 hours resulted in p110α degradation at clinically relevant concentrations (Figure S4A).

To generate direct evidence that taselisib or GDC-0077 treatment was preferentially depleting mutant p110α protein within a mixed WT and mutant allelic population, we took advantage of a neo-tryptic peptide encoded by the *H1047R* mutation to assess p110α protein levels by mass spectrometry. HCC1954 parental cells expressing both mutant and WT p110α were treated with taselisib for 24 hours, revealing loss of the mutant specific peptide only in the taselisib-treated sample, p=0.00016 (Figure 3B). There was no significant change in WT p110α, while the total p110α pool decreased commensurate with the initial abundance of the mutant protein (Figure S4B). Similarly, depletion of the E545K mutant protein was observed in HCC202 heterozygous cells, as shown by assaying a second mutant specific neo-tryptic peptide at this locus, p=0.032 (Figure S4C).

To further explore the molecular mechanisms underlying inhibitor-induced mutant p110α depletion, we examined whether reduction of p110α could be attributed to degradation. A proteasome inhibitor, MG132, and a ubiquitin-activating enzyme E1 inhibitor both rescued the p110α degradation induced by taselisib and GDC-0077 (Figure 3C), while lysosomotropic agents failed to do so (Figure S4D). We next employed ubiquitin pull down assays to confirm that mutant p110α is inducibly ubiquitinated upon taselisib treatment, and that this signal further accumulated when cells were co-treated with MG132 to prevent proteasomal degradation of ubiquitinated p110α (Figure 3D). Accordingly, no ubiquitinated p110α was detected in cells treated with UAE1 inhibitor (Figure 3D). Importantly, taselisib also induced ubiquitination and degradation of E545K p110α in the HCC202 cell model, with comparable rescue by MG132 and UAE1 inhibition (Figure 3D).

Visualization of p110α depletion on western blot was more easily discerned in HCC1954 cells, which have an increased copy number of the mutant *p110α* allele. Quantitative Reverse Transcription PCR (qRT-PCR) analysis confirmed higher expression of the mutant allele (Figure S4E). In order to generate a clean system to better compare differential effect of inhibitors on WT and mutant p110alpha, we used CRISPR/cas9 to generate isogenic HCC1954 cell lines bearing either the *H1047* WT allele or the mutant *H1047R* allele, named HCC1954_mutant and HCC1954_WT (Figure S4E). Taselisib treatment resulted in ubiquitination and depletion of p110α protein only in the HCC1954_mutant cells, and not in the matched HCC1954_WT cells (Figure 3E and Figure S4F).

Since the basal level of ubiquitination was significantly higher for membrane-bound mutant p110α compared with WT p110α (Figure S4G), we hypothesized that the mutant p110α protein may be inherently less stable than the WT protein, as previously noted for mutant EGFR (Greig et al., 2015). Consistent with prior literature (Yu et al., 1998), pulse-chase experiments in HCC1954_WT isogenic cells indicated a protein half-life for WT p110α of ∼26.7 hours. In contrast, the H1047R mutant protein half-life was ∼9.6 hours in the basal state, and was further shortened to ∼4 hours upon treatment with taselisib (Figure 3F). Together these data demonstrate that mutant p110α oncoprotein is inherently less stable and more vulnerable to inhibitor-mediated degradation in a ubiquitin and proteasome dependent manner.

### p85β potentiates p110**α** degradation by recruiting p110**α** to the membrane

It has previously been shown that p110α/p85 dimers are activated by growth factor-stimulated RTK signaling (Cantley et al., 1991). The mechanism of this activation involves release of inhibitory contacts between p110 and p85 once bound to phosphotyrosine residues of an RTK (Backer, 2010; Burke and Williams, 2013). We next reasoned that the ability of our small molecule degraders to accelerate the turnover of activated mutant protein might be through appropriation of the p110α activation process. In support of this hypothesis, cell fractionation studies showed that taselisib-inducible ubiquitination of p110α occurred preferentially in the membrane fraction (Figure 4A).

**Figure 4.**
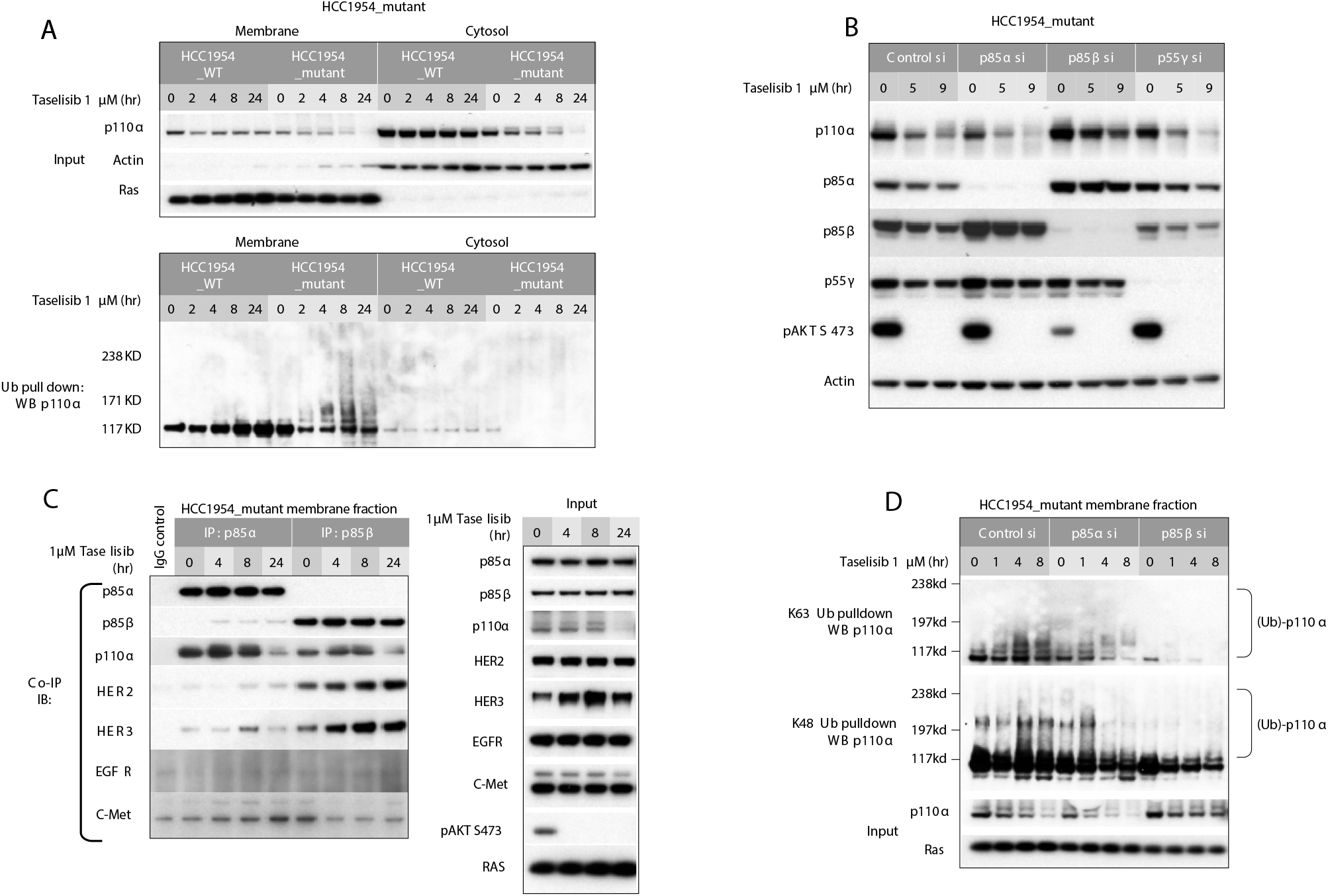
Taselisib-mediated degradation of mutant p110αha occurs preferentially at the plasma membrane. **(A)** Subcellular fractionation of isogenic HCC1954_mutant and HCC1954_wild-type cells. Pull down of ubiquitinated protein was followed by western blotting with anti-p110α. **(B)** HCC1954_mutant cells were transfected with p85α, p85β or p55gγ isoform specific siRNA followed by 1 *µ*M taselisib treatment. **(C)** HCC1954_mutant cell line was treated with 1 *µ*M taselisib for up to 24 hours. Cell lysates were precipitated with p85γ or p85β antibody, followed by immunoblot with antibodies indicated to the left. **(D)** HCC1954_mutant cells were transfected with p85α or p85β siRNA followed by taselisib treatment. Cells were harvested at various time points and fractionated. K63 or K48 linked ubiquitin conjugated protein was pulled down from membrane fraction and analyzed by immunoblotting with p110α antibody.

One major effector that recruits p110α to RTK is the p85 PI3K regulatory subunit. There are three p85 isoforms in class 1A PI3K: p85α, p85β, and p55γ. We first confirmed comparable expression levels of all three isoforms in HCC1954 cells (Figure S5A). By co-immunoprecipitation we also demonstrated that all three isoforms interact with mutant p110α (Figure S5A). We next evaluated the effect of p85 isoforms on inhibitor-induced p110α degradation. Knockdown of both p85α and p55γ showed similar levels of taselisib-induced p110α degradation compared with control siRNA-treated cells. In contrast, we observed that p85β KD rescued taselisib-mediated p110α degradation (Figure 4B). We further observed that p85β KD resulted in inhibition of pathway signaling, shown by reduced pAKT levels. These data suggest that p85 isoforms do not have redundant roles and that the p85β, but not the p85α or p55γ isoforms is involved in mutant p110α degradation. Reduced pAKT levels with p85β KD is most likely a result of reduced p110α membrane localization.

This observation also suggested that p85 isoforms have differential affinity for RTK interaction. To address this question, we next performed a series of co-immunoprecipitation experiments. p85α or p85β were immunoprecipitated from the membrane fraction and immunoblotted with several antibodies, including four receptors that are highly activated in HCC1954 cells (Figure S5B). Both p85α and p85β interacted with p110α, consistent with previous results. We also observed a stronger association between p85β and HER2 and HER3 than observed for p85α or p55γ protein (Figure 4C, Figure S5C). We could not detect an interaction between p85 isoforms with EGFR or c-MET under these conditions.

To further confirm this observation, we next tested whether inhibition of HER2 phosphorylation blocked p85β binding to HER2 and rescue p110α degradation. Cells were treated with taselisib or lapatinib alone or in combination for various timepoints. These cell lysates were next immunoprecipitated using p85α or p85β antibody and immunoblotted with HER2 and HER3 antibodies. In taselisib-treated cells, strong interaction between p85β and HER2/3 was observed. Treatment with lapatinib blocked this interaction, confirming that activated HER2 contributes to p85β membrane binding (Figure S5D). These results further demonstrated that p85β plays an important role in recruiting p110α to the membrane, and that this is likely through binding to activated HER2 and HER3.

To further confirm the role of p85β in p110α degradation, we next tested the effects of p85 knockdown on K63 and K48 polyubiquitin chain formation on p110α. Upon knockdown of p85α or p85β and following inhibitor treatment, K63 or K48 ubiquitin conjugated proteins were immunoprecipitated using linkage specific antibodies and immunoblotted with p110α. In both control- and p85α-depleted cells, taselisib treatment induced both K63 and K48 linked ubiquitination on p110α. However, upon p85β depletion, ubiquitination of p110α was no longer detected (Figure 4D). These data further confirm that the p85β regulatory subunit plays an important role in taselisib-induced mutant p110α degradation, by recruiting p110α to the membrane where ubiquitination occurs.

### Taselisib- and GDC-0077-induced mutant p110α degradation is dependent on RTK activity

To further understand whether inhibitor-induced mutant p110α degradation is dependent on its recruitment to activated RTK, we next investigated the efficacy of the two clinically relevant molecules, GDC-0077 and BYL719 (Alpelisib). Unexpectedly, we observed a significant difference in the sensitivity of *HER2*-amplified (∼20-fold difference between the mean IC^50^ values) versus *HER2*-negative cell lines (∼6-fold difference between two inhibitors) to GDC-0077 versus BYL719. The inhibitors were not differentiated in WT cell lines regardless of *HER2* status (Figure 5A). These data imply that RTK activity may define the sensitivity of a cell line to inducible degradation of mutant-p110α. To confirm this finding, a panel of over 50 cell lines harbouring *PIK3CA* hot spot mutations were analyzed (Figure 5B). Most of these were heterozygous, carrying both WT and mutant *PIK3CA* alleles at differing frequencies. For easier visualization of mutant p110α depletion, we analyzed the cell lines with higher copy numbers of mutant alleles that also represented each hot spot mutation across various tumors (Figure 5B, colored in blue). In a cell proliferation assay, these cell lines have varying GDC-0077 sensitivity. All of the selected representative cell lines responded to GDC-0077, as measured by inhibition of pAKT. Not all cell lines showed visible p110α degradation; those that showed mutant p110α degradation were HCC2185, HCC1954, MDAMB453, and KPL4 (Figure 5B). It was particularly striking that all *HER2*-amplified cell lines showed p110α degradation. In contrast, the *HER2*-negative cell lines were resistant to degradation except HCC2185. To further understand this observation, we tested if activating RTK by addition of growth factors would induce p110α degradation. A subset of *HER2*-negative cell lines, each harboring WT or one of three p110α hot spot mutations, were treated with GDC-0077 alone or in combination with growth factors. All responded to growth factors, as shown by induction of HER3 phosphorylation. Consistent with our data, p110α levels did not decrease in *HER2*-negative cell lines treated with GDC-0077 alone. However, the combination of growth factors and GDC-0077 induced p110α degradation in all three p110α-mutant lines. WT p110α protein expression was not affected by addition of growth factors (Figure 5C). Immunoprecipitation and pull down of p110α revealed that p110α degradation was markedly delayed when RTK interaction with the p110α/p85 complex was disrupted, with limited inducible degradation observed at the 8 hr timepoint (Figure 5D, upper panel). Conversely, inhibition of RTK phosphorylation by lapatinib partially rescued taselisib-mediated p110α degradation in HCC1954 cells (Figure 5D, lower panel). Similar to taselisib, GDC-0077-induced mutant p110α degradation was rescued by lapatinib treatment (Figure 5E). In support of these results, we observed that in multiple patient derived xenograft (PDX) models, high basal pHER2 and pHER3 expression level correlated with better taselisib-mediated p110α degradation, with weaker degradation observed in tumors with low levels of pRTK (Figure S6A). In addition, within this group of PDX models, the degree of degradation appeared to correlate with basal pRTK expression. Taken together, our data support a model where small molecule-induced p110α mutant degradation depends on RTK activity in *PIK3CA* mutant cancers.

**Figure 5.**
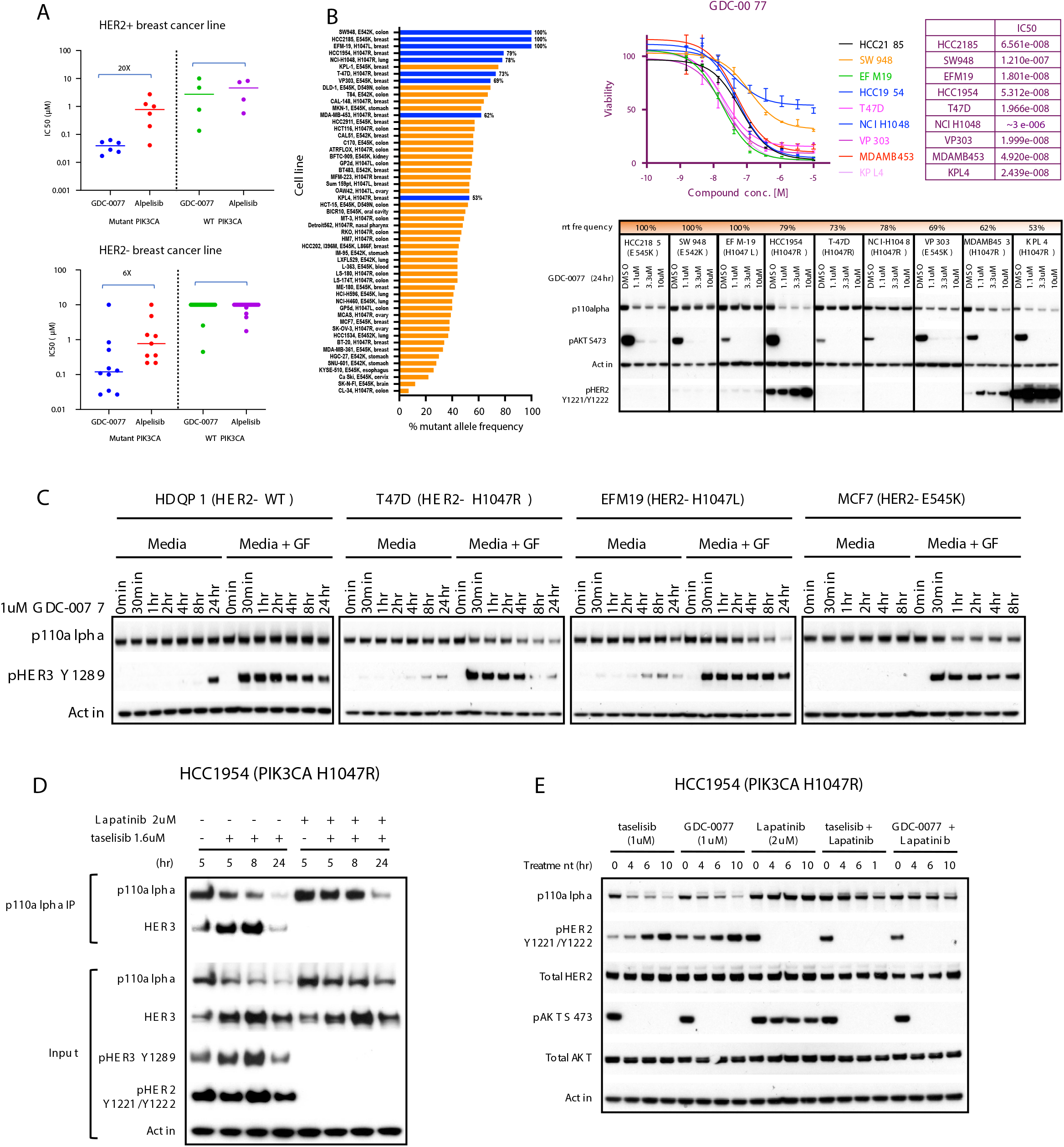
Taselisib- and GDC-0077-induced mutant p110α degradation is dependent on RTK activity. **(A)** Cell viability IC_50_ values determined by quantifying ATP from breast tumor lines: HER2-positive *PIK3CA*-mutant (n=6), HER2-positive *PIK3CA*-wild-type (n=4), HER2-negative *PIK3CA*-mutant (n=10), and HER2-negative *PIK3CA*-wild-type (n=20) at 5 days post-treatment. **(B)** Bar plot showing *PIK3CA*-mutant frequency among tumor lines harboring *PIK3CA* hotspot mutations. ATP based cell viability assay in selected cell line (HCC2185, SW948, EFM-19, HCC1954, T-47D, NCI-1048, VP303, MDAMB453 and KPL4). western blot of the p110α protein levels and pAKT signaling in same cell lines treated with GDC-0077 for indicated concentrations for 24 hours. **(C)** HER2-negative *PIK3CA*-wild-type or -mutant cells cultured in standard media with 10% FBS and treated with GDC-0077 alone or addition of growth factors (EGF and neuregulin). **(D)** HCC1954 PIK3CA H1047R cells treated with taselisib alone or combination of taselisib and lapatinib. Cell lysates were precipitated with p110α antibody, followed by western blot with HER3 antibody. **(E)** Cell lysates following treatment with taselisib or GDC-0077 alone or combination with lapatinib for indicated time points followed by western blot analysis with indicated antibodies.

### p110**α**-mutant degrading inhibitors provide more sustained benefit in HER2-positive versus HER2-negative p110**α**-mutant cancers

Given that RTK reactivation could potentially limit efficacy, we also aimed to understand activity in HER2-positive breast cancers. Degradation of mutant p110α blocked the feedback induced pathway reactivation and resulted in enhanced potency of taselisib and GDC-0077 in cellular assays and an increase in apoptosis in mutant cells. Our data thus far predict that drugs which induce degradation of mutant p110α may show more sustained benefit over non-degraders by preventing pathway reactivation. We speculated that the extent of negative feedback depends on the amount of RTK expression, implying that inhibitor-mediated feedback pathway reactivation would be weak in *HER2*-negative cells. Also, based on our findings, low RTK activity may result in inefficient mutant p110α degradation.Pathway reactivation was not observed in *HER2*-negative cells, in contrast to *HER2*-amplified cells, irrespective of the inhibitors used. This suggests that in *HER2*-negative mutant cells, the degrader mechanism of action may not provide additional benefit over drugs with a non-degrader mechanism (model in Figure 6A). This was further confirmed in a panel of *PIK3CA* mutant cell lines. In *HER2*-amplified cells, GDC-0077 resulted in sustained pathway inhibition, while BYL719 activity was attenuated, as evidenced by rebound of pAKT levels (Figure S6B). In contrast, in a panel of *HER2*-negative lines, there was no difference in pathway inhibition by both inhibitors, GDC-0077 or BYL719.

**Figure 6:**
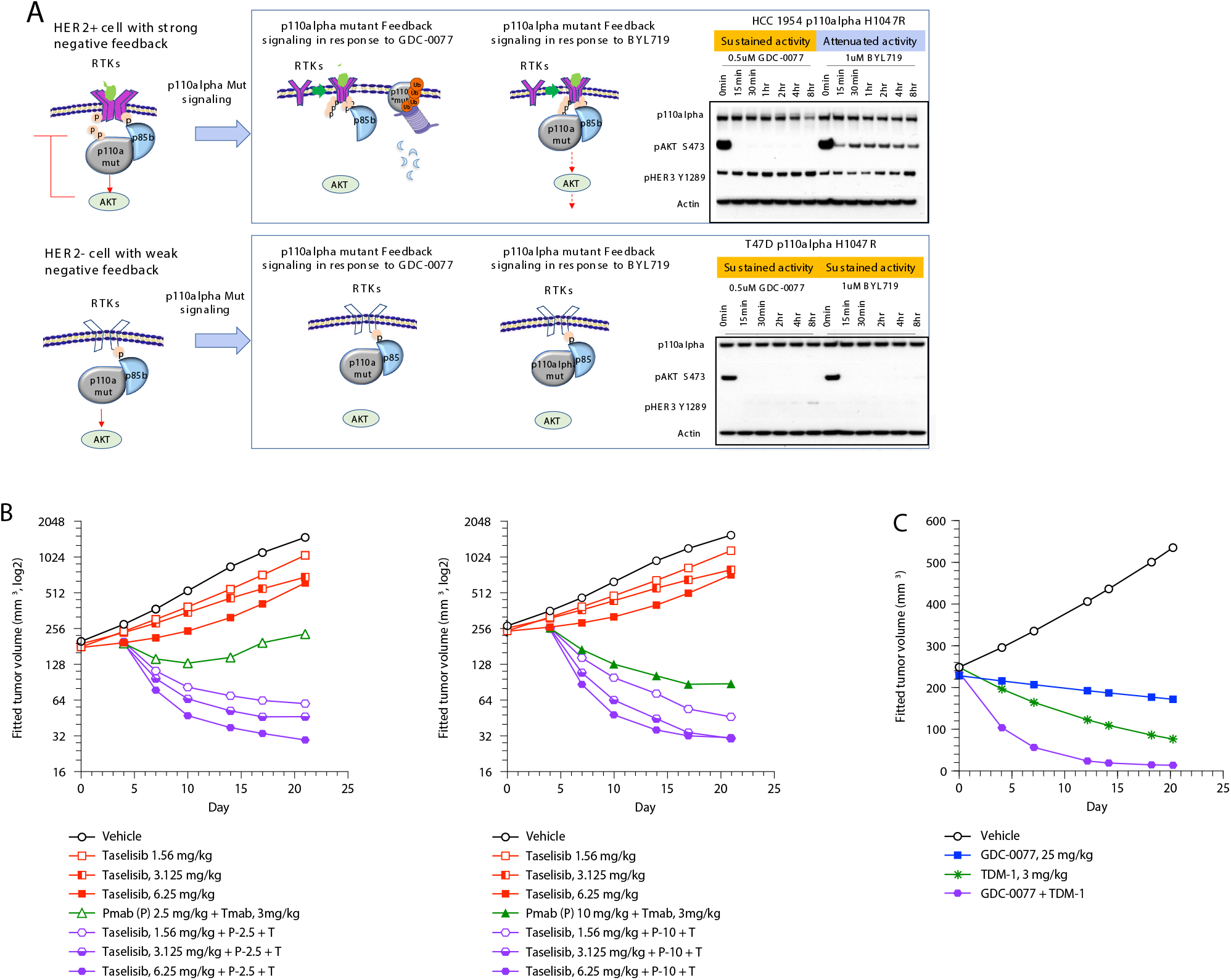
p110α-mutant degrading inhibitors provide more benefit in HER2-positive versus HER2-negative p110α-mutant cancers. **(A)** Mechanistic model of drug-induced p110α degradation in HER2-positive and HER2-negative *PIK3CA*-mutant cells. **(B)** Tumor growth curve from KPL-4 (HER2+, PIK3CA H1047R) xenograft treated with vehicle, taselisib, trastuzumab plus pertuzumab, or the indicated drug combination. **(C)** Tumor growth curve from KPL-4 (HER2+, PIK3CA H1047R) xenograft treated with vehicle, GDC-0077, TDM-1 or the indicated drug combination.

Based on these data, we posited that we could best leverage the degradation potential of GDC-0077 in *HER2*-amplified cancers. In HER2-driven breast cancers, HER2-targeted therapy is the standard of care. However, *HER2*-amplified tumors with p110α mutations are less responsive to HER2-targeted therapy. Therefore, combination treatment of the PI3K pathway inhibitors and HER2 targeted therapy should result in enhanced efficacy. Indeed, combination of HER2 inhibitors, trastuzumab and pertuzumab, in combination with taselisib in a HER2-positive mutant p110α KPL-4 xenograft model showed better response compared with single drugs alone (Figure 6B. Furthermore, combining ado-trastuzumab emtansine (TDM-1) with GDC-0077 showed a synergistic effect in the KPL-4 xenograft model as well (Figure 6C).

Given that most p110α-mutant cells are heterozygous, selective mutant p110α degradation would imply a possible scenario where WT p110α protein is still present and can reactivate the same pathway and dampen inhibitor activity (Figure S6C). To test this possibility, we compared levels of feedback reactivation in an engineered homozygous versus heterozygous line. In a HCC1954 WT homozygous isogenic line, the pathway was reactivated by both GDC-0077 (degrader) and BYL719 (non-degrader), as expected (Figure S6D). In a HCC1954 mutant homozygous isogenic line, BYL719, but not GDC-0077 treatment reactivated pathway signaling. Importantly, when compared, the phenotype of a heterozygous parental line behaved similar to that of a p110α mutant homozygous line, suggesting that in heterozygous cells, mutant p110α functions as the main driver of cell signaling (Figure S6D). Furthermore, cell proliferation assays mirrored the inhibitor efficacy as well. GDC-0077 showed the same efficacy between homozygous and heterozygous mutant lines but showed reduced efficacy in WT isogenic lines (Figure S6D), while no shift in IC_50_ was observed between all lines following BYL719 treatment. To further confirm these results, the same experiment was performed in *HER2*-negative lines. Consistent with our hypothesis for the role of HER2 in pathway reactivation, feedback pathway reactivation was not detected even in a homozygous WT line upon treatment with GDC-0077 or BYL719. In cell viability assays, neither inhibitor showed a shift in efficacy between isogenic lines (Figure S6E). Taken together, our data demonstrate that degradation potential of GDC-0077 would be most leveraged in HER2-positive breast cancers for more sustained pathway inhibition.

## Discussion

In summary, we have discovered that the mutant p110α oncoprotein has unique characteristics compared with WT p110α: a shorter half-life, ubiquitination in the membrane fraction, and proteasome-mediated turnover. Furthermore, the mutant oncoprotein is susceptible to increased proteasome-mediated degradation upon binding particular PI3K inhibitors such as taselisib and GDC-0077. Our results suggest that RTK activity plays a key role in regulating p110α degradation by recruiting p110α to the membrane. The mutant oncoprotein may be particularly vulnerable to additional local conformational changes that impact membrane binding, and taselisib and GDC-0077 may be enhancing this effect to accelerate proteasome-mediated degradation. This discovery reveals a new mechanism of action to exploit in *PIK3CA*-mutant tumors, opening an exciting path to increasing drug efficacy for the predominant oncogene in cancer. With a combined mutant-p110α degrader mechanism and exquisite p110α isoform selectivity, the PI3Kα-selective inhibitor and mutant PI3Kα degrader GDC-0077 may provide opportunities for previously inaccessible combination therapy with greater clinical responses. Moreover, this suggests that it may be possible to exploit endogenous mechanisms and the intrinsic instability of mutant oncoproteins more broadly to develop tumor-selective therapeutic agents. Based on these mechanistic discoveries, it may be possible to create efficacious compounds that specifically deplete the mutant p110α protein without blocking WT PI3K signaling, similar to engineered mouse models of cancer in which removal of the mutant *PIK3CA* oncogene induces regressions in the presence of WT PI3K (Cheng et al., 2016; Engelman et al., 2008). For the over 2 million cancer patients diagnosed annually with *PIK3CA*-mutant tumors, this discovery opens the possibility of a future therapeutic agent that solely targets tumor cells bearing mutant *p110*α without the systemic adverse effects of inhibiting WT signaling.

With respect to the clinical relevance of our findings, a first-in-human, open-label, Phase I/IB dose escalation study of oral daily GDC-0077 alone and in combination with endocrine and targeted therapies for *PIK3CA*-mutant solid tumors is ongoing. The single-agent portion of the study showed that GDC-0077 had a manageable safety profile with a maximum tolerated dose of 9 mg once daily (supported by a linear pharmacokinetic profile), with promising anti-tumor activity (Juric et al., 2019). When combined at this dose with letrozole with and without palbociclib (Jhaveri et al., 2019), or when combined with fulvestrant the safety profile was also manageable with promising anti-tumor activity (Kalinsky et al., 2020). No drug-drug interactions were observed in the letrozole ± palbociclib portion of the study (Jhaveri et al., 2019). Phase III trials are now ongoing to assess efficacy and safety in a randomized, controlled manner in locally advanced or metastatic HR-positive/HER2-negative breast cancer in combination with palbociclib and fulvestrant (NCT04191499). To further leverage the degradation potential of GDC-0077 in HER2-positive breast cancers, this work has also provided the rationale for a first-in-human, open-label, Phase I/IB dose escalation study of oral daily GDC-0077 in combination with standards of care (trastuzumab and pertuzumab) is also being enabled.

## Methods

### Chemical reagents

The proteasome inhibitor MG132 (474790) was obtained from Calbiochem EMD Millipore (Billerica, MA). Chloroquine (14774) was obtained from Cell Signaling (Danvers, MA). Ammonium Chloride NH4Cl (254134) was obtained from Sigma-Aldrich (St. Louis, MO). PI3K inhibitors and ubiquitin-activating E1 inhibitor (Chen et al., 2011) were provided by the chemistry department at Genentech, Inc. (South San Francisco, CA).

### Antibody reagents

Antibodies to p110α (4249), cleaved poly (ADP-ribose) polymerase (PARP) (9541), phosphorylated (phospho)-Akt Ser473 (4060), pS6 S235/236 (2111), anti-ubiquitin (3936), p110δ (34050), HER3 (12708), HER2 (2242), pHER2 Y1221/Y1222 (2243), and pHER3 Y128 (4791) were obtained from Cell Signaling (Danvers, MA). The antibody to β-actin (A5441) was from Sigma-Aldrich. Antibodies to Ras (ab52939), p85α (ab133595), and p85β (ab28356) were obtained from Abcam (Cambridge, MA). Ubiquitin reagent TUBE1 (UM101) was obtained from LifeSensors, Inc. (Malvern, PA). The p55γ antibody (MAB6638) was obtained from R&D Systems (Minneapolis MN). Antibodies to GAPDH (MAB374) and p110β (04-400) were obtained from EMD Millipore.

### pPRAS40 ELISA assay

SW48 isogenic cells were plated in 384-well tissue-culture treated assay plates (Cat. No. 781091; Greiner Bio-One; Monroe, NC) and incubated overnight at 37°C and 5% CO_2_. The three isogenic SW48 lines (WT, E545K, and H1047R) were plated and assayed in parallel. The following day, test compounds were serially diluted in DMSO and added to cells (final DMSO concentration of 0.5%). Cells were then incubated with drugs for 24 hours at 37°C and 5% CO_2_. After 24 hours, cells were lysed and phosphorylated proline-rich AKT substrate of 40 kDa (pPRAS40) levels were measured using the Meso Scale Discovery (MSD^®^) custom pPRAS40 384w Assay Kit (Cat. No. L21CA-1; Rockville, MD). Cell lysates were added to assay plates pre-coated with antibodies against pPRAS40 and allowed to bind to the capture antibodies overnight at 4°C. The detection antibody (anti-total pPRAS40, labeled with an electrochemiluminescent SULFO-TAG^TM^) was added to the bound lysate and incubated for 1 hour at room temperature. The MSD^®^ Read Buffer was added such that when voltage was applied to the plate electrodes, the labels bound to the electrode surface emitted light. The MSD^®^ Sector Instrument measured the intensity of the light and quantitatively measured the amount of pPRAS40 in the sample. Percent inhibition of pPRAS40 per concentration of compounds was calculated relative to untreated controls. The EC_50_ values were calculated using the 4-parameter logistic nonlinear regression dose-response model. Reported EC_50_ values indicate an average value from three independent experiments. Standard deviations are reported as ± the reported EC_50_ values for each cell line. The EC_50_ in WT cells was divided by EC_50_ in mutants to derive the fold-increase in potency in mutant cells.

### Viability assay CellTiter-Glo^®^

Cells were seeded (1000–2000 cells/well) in 384-well plates for 16 hours. On day two, nine serial 1:3 compound dilutions were made in DMSO in a 96-well plate. The compounds were then further diluted into growth media using a Rapidplate robot (Zymark Corp., Hopkinton, MA). The diluted compounds were then added to quadruplicate wells in the 384-well cell plate and incubated at 37°C and 5% CO_2_. After 4 days, relative numbers of viable cells were measured by luminescence using CellTiter-Glo® (Promega) according to the manufacturer’s instructions and read on a Wallac Multilabel Reader (PerkinElmer, Foster City). The EC^50^ calculations were carried out using Prism 6.0 software (GraphPad, San Diego). The GR calculations and figures were performed using R scripts based on Hafner et al. (Hafner et al., 2016).

### Nucleosome ELISA

MDA-MB-453, HCC202, and Cal85-1 cells were plated in 96-well tissue-culture treated assay plates (Corning; Cat. No. 3904; Corning, NY) and incubated overnight at 37°C and 5% CO_2_. The following day, PI3K inhibitors were serially diluted in DMSO and added to cells (final DMSO concentration of 0.5%). Cells were then incubated with drugs for 72 hours at 37°C and 5% CO_2_. After 72 hours, cells were lysed and centrifuged at 200 g for 10 min. Histone-associated DNA-fragment levels were analyzed using the Cell Death Detection ELISAPLUS (Roche; Cat. No. 11920685001; Basel Switzerland). 20 *µ*L of supernatant was added to each well of the streptavidin capture plate followed by 80 *µ*l of anti-histone-biotin, and DNA-peroxidase immunoreagent. Plates were incubated at room temperature for 2 hours shaking (300 RPM). Contents of the plate were removed followed by three washes with Incubation buffer. 100 *µ*L of ABTS solutions was added to each well and plates were incubated for 10–20 min, after which 100 *µ*L of ABTS Stop Solution was added to each well. Absorbance was measured at 405 nm and 490 nm of each plate. Fold increases were generated by assessing the increased of absorbance (A405 nm–A490 nm) of wells from cells treated with compounds, normalized to those treated with DMSO alone.

### Cell lines and cell culture

Cell lines were obtained from the ATCC. All cell lines underwent authentication by Short Tandem Repeat profiling, SNP fingerprinting, and mycoplasma testing (Yu et al., 2015). The isogenic colon cancer cell lines SW48 human PIK3CA (H1047R/+) (HD103-005) and SW48 human PIK3CA (E545K/+) (HD103-001) and SW48 parental line were obtained from Horizon Discovery Ltd. (Cambridge, UK). Cell lines were grown under standard tissue-culture conditions in RPMI media with 10% fetal bovine serum (Gibco, 10082-147), 100 U/mL penicillin-streptomycin (Gibco, 15140-122), 2 mmol/L L-glutamine (Gibco, 15030-081). *PIK3CA* mutation status and frequencies of all cell lines are summarized in Figure 5B. Cells were treated with compounds for the indicated periods of time. For rescue experiments, 1 *µ*M final concentration of proteasome inhibitor MG132 (EMD Millipore; Cat. No. 474790-5MG; Darmstadt, Germany), 2 *µ*M HER2 inhibitor lapatinib (Selleckchem; Cat. No. GW-572016), or 2 *µ*M Ubiquitin activating enzyme E1 inhibitor (synthesized at Genentech), were added before cell harvest.

### CRISPR engineering of HCC1954 cells

HCC1954 breast cancer cells were engineered using CRISPR to knock out either the *PIK3CA* WT allele or the mutant alleles, to create an isogenic pair of cell lines designated as HCC1954_mutant and HCC1954_WT. gRNAs were designed using the CRISPRtool (http://crispr.mit.edu) to minimize potential off-target effects and cloned into pCas-Guide-EF1a-GFP vector (Blueheron Biotech). To generate the HCC1954_mutant line bearing p110α H1047R as homozygous mutant, two CRISPR-Cas9 constructs were designed. One was designed to specifically target the wild type allele in exon 21 (gRNA H1047R-2, ATGAATGATGCACATCATGG) and the second designed to target the intron of both WT and mutant alleles (gRNA H1047R-7, ACATTTGAGCAAAGACCTGA). To generate the HCC1954_WT line, two CRISPR-Cas9 constructs were designed with one gRNA targeting the mutant allele in exon 21 (H1047R-5, ATGAATGATGCACGTCATGG) and the second gRNA targeting the intron of both WT and mutant alleles (H1047R-8, TATTAAACTCCTGACATGCC). Plasmids for each targeting pair were co-transfected using Turbofectin (Thermofisher). After 48 hours cells were put under selection with 1 ug/mL puromycin. Puromycin resistant cells were further selected by collecting GFP expressing cells by flow cytometry, and clones were expanded in standard cell culture conditions to create stable lines. Targeting efficiency of the CRISPR-induced allelic knockouts was assessed by PCR flanking the target sites (forward: TGCTGTGAAGGAAAATGGAA, (reverse: TGCAGTGTGGAATCCAGAGTGAGC), and clones further validated with qRT-PCR and DNA sequencing.

### siRNA transfection

Transfection of siRNA was carried out using Lipofectamine RANiMAX reagent (Thermofisher), 72 hours in advance of drug treatment.

### Western blots

Protein concentration was determined using the Pierce BCA Protein Assay Kit. For immunoblots, equal protein amounts were loaded and then separated by electrophoresis through NuPAGE Novex Bis-Tris 4–12% gradient gels (Invitrogen). Proteins were transferred onto Nitrocellulose membranes using the iBlot system and protocol from InVitrogen (Carlsbad, CA).

### Subcellular fractionation

Cells were washed once with phosphate-buffered saline, before scraping into 0.8 ml/dish hypotonic lysis buffer (HLB: 25 mM Tris–HCl pH7.5, 10 mM NaCl, 1 mM EDTA, protease and phosphatase inhibitors). The cells were lysed by 30 strokes in a Dounce homogenizer, subjected to centrifugation at 1500 g (3000 RPM) for 5 min to pellet nuclei and unbroken cells, followed by centrifugation of the supernatant at 100,000 g (44,000 RPM) in TLA55 rotor for 40 min. The supernatant (800 *µ*L) was collected (S100 fraction) and the pellet resuspended in 200ul HLB plus 1% NP40 (P100 fraction). The resuspended pellet was centrifuged 5 min in high speed in microfuge and supernatant collected.

### Immunoprecipitation and pulldown

Cells were lysed in 20 mM TrisHCL pH 7.5, 137 mM NaCl, 1 mM EDTA, 1% NP40, 10% glycerol plus protease and phosphatase inhibitors. For p110α immunoprecipitation, lysates were incubated with p110α antibody (Cell Signaling, 4249) overnight. 50 *µ*L of proteinA agarose beads were added to each sample and incubated additional 2 hours. For ubiquitinated protein pulldown experiment, cells were lysed in lysis buffer containing 200 *µ*g/mL TUBE1 (Lifesensors UM101). Lysates were isolated and added 50 *µ*L of glutathione agarose beads (Sigma, G4705). The samples were incubated overnight and captured ubiquitinated protein was eluted in SDS reducing sample buffer.

### RNA isolation and *PIK3CA* allele-specific quantitative RT-PCR

Total RNA was isolated from cells using RNeasy Plus Mini Kit (Qiagen) following the protocol described in the kit. First-strand cDNA synthesis and RT-qPCR was carried out using One step RT QPCR reagent (Roche). Resulting signal was detected on Applied Biosystems Real Time PCR System. Primers and allele specific probes were:

*PIK3CA* H1047R-forward: GGCTTTGGAGTATTTCATGAAACA

*PIK3CA* H1047R-reverse: GAAGATCCAATCCATTTTTGTTGTC

*PIK3CA* H1047R WT-probe: ATGATGCACATCATGGT

*PIK3CA* H1047R Mut-probe: TGATGCACGTCATGGT

*PIK3CA* E545K-forward: GCAATTTCTACACGAGATCCTCTCT

*PIK3CA* E545K-reverse: CATTTTAGCACTTACCTGTGACTCCAT

*PIK3CA* E545K WT-probe: TGAAATCACTGAGCAGGAG

*PIK3CA* E545K Mut-probe: TGAAATCACTAAGCAGGA

### Intracellular drug concentration

Taselisib (1 *µ*M in RPMI medium) was added to *PIK3CA*-mutant and -WT cancer cell lines in triplicate and incubated under standard culture conditions (37°C, 5% CO_2_) for 18 hours. Medium containing taselisib was aspirated and cells were washed twice with 1 mL of ice-cold Hank’s balanced salt solution (HBSS). Cells were lysed by sonication for 1 min in HBSS followed by the addition of an equal volume of acetonitrile containing analytical internal standard (propranolol, 100 nM). The supernatant was analyzed in positive mode on a SCIEX API-4000 LC/MS/MS system with the transitions of 461/418 and 260/116 for taselisib and propranolol, respectively. The LC separations were achieved using a LUNA C18 column (4 *µ*m, 50 x 2.1 mm) from Phenomenex, Inc. (Torrance, CA) and Agilent 1100 series pumps consisting of mobile phase A (water containing 0.1% formic acid) and mobile phase B (acetonitrile containing 0.1% formic acid). The flow rate through the system was 1 mL/min. The initial condition was set at 5% B, which was ramped to 95% B over 40 sec. After allowing the system to hold at 95% B for 20 sec, the gradient was changed back to the initial condition of 5% B and was allowed to equilibrate for 54 sec before the next injection. A parallel replicate plate containing *PIK3CA*-mutant and -WT cancer cells was reserved for the determination of protein concentrations. Cells were washed with HBSS buffer and then lysed with 0.5 mL of ProteaPrep cell lysis solution (Protea Biosciences, Morgantown, WV). After 30 min, the solution was neutralized with 0.5 mL HBSS buffer and 100 μL of the lysed cells were used to determine the protein concentration by the standard Bradford protein assay (Thermo Scientific, Rockford, IL), with bovine serum albumin as a standard.

### PI3K ADP-Glo assays for Ki measurement

As described in WO 2017/001645 A1, the PI3K lipid kinase reaction was performed in the presence of PIP2:3PS lipid substrate (Promega #V1792) and ATP. Following the termination of the kinase reaction, turnover of ATP to ADP by the phosphorylation of the lipid substrate was detected using the Promega ADP-Glo™ (Promega #V1792) assay. Reactions were carried out using the following conditions for each PI3K isoform: PI3Kalpha (Millipore #14-602-K) kinase concentration 0.2 nM, ATP 40 *µ*M, PIP2:3PS 50 *µ*M; PI3Kbeta (Promega #V1751) kinase concentration 0.6 nM, ATP 40 *µ*M, PIP2:3PS 50 *µ*M; PI3K delta (Millipore #14-604-K), kinase concentration 0.25 nM, ATP 40 *µ*M, PIP2:3PS 50 *µ*M; PI3K gamma (Millipore #14-558-K), kinase concentration 0.4 nM, ATP 25 *µ*M, PIP2:3PS 50 *µ*M. After 120 min of reaction time, the kinase reaction was terminated. After the reaction, any ATP remaining was depleted leaving only ADP. Then the Kinase Detection Reagent was added to convert ADP to ATP, which was used in a coupled luciferin/luciferase reaction. The luminescent output was measured and was correlated with kinase activity. All reactions were carried out at room temperature. For each PI3K isoform a 3 *µ*L mixture (1:1) of enzyme/lipid substrate solution was added to a 384 well white assay plate (Perkin Elmer #6007299) containing 50 *µ*L of test compound or DMSO only for untreated controls. The reaction was started by the addition of 2 *µ*L ATP/MgCl_2_. The kinase reaction buffer contained 50 mM HEPES, 50 mM NaCl_2_, 3 mM MgCl_2_, 0.01% BSA, 1% DMSO, and enzyme and substrate concentrations. The reaction was stopped by the addition of 10 *µ*L ADP-Glo reagent. Plates were read in a Perkin Elmer Envision system using luminescence mode. A 10-point dose response curve was generated for each test compound. Ki values for each compound were determined using the Morrison Equation.

### Kinetic solubility

Compounds were dissolved in DMSO to a concentration of 10 mM. These solutions were diluted into PBS buffer (pH 7.2, composed with NaCl, KCl, Na_2_HPO_4_, and KH_2_PO_4_) to a final compound concentration of 100 μM, DMSO concentration of 2%, at pH 7.4. The samples were shaken for 24 hours at room temperature followed by filtration. LC/CLND was used to determine compound concentration in the filtrate, with the concentration calculated by a caffeine calibration curve and the samples nitrogen content. An internal standard compound was spiked into each sample for accurate quantification.

### Plasma protein binding

As described in literature (Heffron et al., 2016), the extent of protein binding was determined in vitro, in CD-1 mouse, Sprague−Dawley rat, and human plasma (Bioreclamation, Inc., Hicksville, NY) by equilibrium dialysis using the RED Device (Thermo Fisher Scientific, Rockford, IL). Compounds were added to pooled plasma (n ≥ 3) at a total concentration of 5 μM. Plasma samples were equilibrated with phosphate-buffered saline (pH 7.4) at 37°C in 90% humidity and 5% CO_2_ for 4 hours. Following dialysis, compound concentration in plasma and buffer were measured by LCMS/MS. The percent of unbound compound in plasma was determined by dividing the concentration measured in the post-dialysis buffer by that measured in the post-dialysis plasma and multiplying by 100.

### MDCK permeability

Experiments were carried out as previously described (Irvine et al., 1999).

### Drug half-life *in vivo*

Experiments to determine half-life in mice were carried out as previously described (Furet et al., 2013; Pang et al., 2014; Salphati et al., 2011).

### Pulse-chase

Cells were starved in methionine/cysteine deficient media overnight. Cells were pulse labeled with 35S cysteine and methionine (Pro-mix l-[35S], Amersham) in RPMI lacking cysteine and methionine for 6 hours, followed by three washes with RPMI containing no label. Cells were then incubated in normal media containing methionine and cysteine, either with or without compounds. Lysates were collected at various time points up to 48 hours. Radiolabeled p110α was immunoprecipitated and run on SDS-PAGE. Images were acquired on the Typhoon Scanner (GE Healthcare-Amersham), and signals quantified and normalized using ImageQuant TL software. Data was fit to a single exponential decay function using Prism (GraphPad, San Diego, CA) with Y=(Y0-NS)*exp(-K*X) + NS and constraints Y0=100, NS=0.

### Animal studies

The in vivo efficacy of PI3K inhibitors was tested in three breast cancer xenograft models that harbor *PIK3CA* mutations, HCC1954 (HER2-positive, *PIK3CA* H1047R), WHIM20 (ER-positive/HER2-negative, *PIK3CA* H1047R) and HCI-003 (ER-positive/HER2-negative, *PIK3CA* E545K). All in vivo studies were approved by Genentech and Institutional Animal Care and Use Committee (IACUC) and adhered to the NIH Guidelines for the Care and Use of Laboratory Animals. HCC1954 tumor cells (5×106) were inoculated in the 2/3 mammary fat pads of female NCR nude mice (Taconic Farms, Hudson NY) while WHIM20 and HCI-003 tumors (50mm^3^) were engrafted in 2/3 mammary fat pads in female NOD-SCID gamma mice (Jackson Laboratories, Bar Harbor, Maine). Tumor volumes were measured using Ultra Cal-IV calipers (Model 54-10-111; Fred V.Fowler Co.; Newton, MA). The following formula was used in Excel, version 11.2 to calculate tumor volume: Tumor Volume (mm^3^) = (Length x Width^2^) x 0.5. Mice for efficacy studies were distributed into 8–10 mice/group with a mean tumor volume of 200 to 250 mm^3^ at the initiation of dosing. A linear mixed effect (LME) modeling approach was used to analyze the repeated measurement of tumor volumes from the same animals over time (Pinheiro et al., 2017). Cubic regression splines were used to fit a non-linear profile to the time courses of log^2^ tumor volume at each dose level. These non-linear profiles were then related to dose within the mixed model. Tumor growth inhibition as a percentage of vehicle control (%TGI) was calculated as the percentage of the area under the fitted curve (AUC) for the respective dose group per day in relation to the vehicle, using the following formula: %TGI=100 x (1 - AUCdose/AUCvehicle). The PI3K inhibitors taselisib, GDC-0941, GDC-0077, and BYL719 were formulated in Methylcellulose Tween (MCT) vehicle consisting of 0.5% (w/v) methylcellulose, 0.2% (w/v) polysorbate 80 (Tween-80) and dosed orally by gavage daily. Tumor sizes and mouse body weights were recorded twice weekly, and mice with tumor volume exceeding 2000 mm^3^ or body weight loss of 20% of starting weight were promptly euthanized. For pharmacodynamic (PD) and mechanistic studies, mice were dosed once by oral gavage and tumors harvested at 4 hours post-dose. Following drug treatment, tumors were harvested and snap-frozen in liquid nitrogen, dissociated in lysis buffer containing 10 *µ*M Tris pH 7.4, 100 *µ*M NaCl, 1 *µ*M EDTA, 1 *µ*M EGTA, 1 *µ*M NaF, 20 *µ*M Na_4_P_2_O_7_, 2 mM Na_3_VO_4_, 1% Triton X-100, 10% glycerol, 0.1% SDS, and 0.5% deoxycholate supplemented with a phosphatase and protease inhibitor cocktail (Sigma, St. Louis, MO). Protein concentrations were determined in whole cell lysates using the BCA Protein Assay Kit (Pierce; Rockford, IL). Membrane fractions were isolated from xenograft tumors as described above, and assessed by western blot.

### Mass spectrometry

Liquid chromatography-tandem mass spectrometry (LC-MS/MS) analysis was performed on p110α protein immunoprecipitated from three cell lines: HCC1954, HCC202 and HDQ-P1. Each cell line was treated for 24 hours with either DMSO (vehicle) or taselisib (500 nM). Each experiment was performed beginning with 4–6 mg protein lysate per cell line/treatment (total of 6 samples/experiment) for n=4 biological replicates. One gel region per sample, corresponding to the expected migration was excised based on the migration of purified p110α protein in an adjacent lane. Gel pieces were diced into ∼1 mm^3^ pieces and subjected to in-gel digestion as follows. Gel pieces were de-stained with 50 mM ammonium bicarbonate/50% acetonitrile and dehydrated with 100% acetonitrile prior to reduction and alkylation using 50 mM dithiothreitol (30 min, 50°C) and 50 mM iodoacetamide (20 min, room temperature), respectively. Gel pieces were again dehydrated, allowed to reswell in a 20 ng/*µ*L trypsin in 50 mM ammonium bicarbonate/5% acetonitrile digestion buffer on ice for 2 hours, and then transferred to a 37°C oven for overnight incubation. Digested peptides were transferred to microcentrifuge tubes and gel pieces were extracted twice, once with 50% acetonitrile/0.5% trifluoroacetic acid, and a second round with 100% acetonitrile. Extracts were combined with digested peptides and speed-vac dried to completion. Samples were reconstituted in 5% formic acid/0.1% heptafluorobutyric acid/0.01% hydrogen peroxide 30 min prior to LC-MS/MS analysis. Samples were analyzed by LC-MS/MS with duplicate injection (with the exception of the first replicate where samples were injected once) on a Thermo LTQ Orbitrap Elite coupled to a Waters nanoAcquity UPLC. Peptides were loaded onto a 0.1mm X 100mm Waters Symmetry C18 column packed with 1.7 *µ*m BEH-130 resin and separated via a two-stage linear gradient where solvent B (98% acetonitrile, 2% water) was ramped from 5% to 25% over 20 min and then from 25% to 50% over 2 min. Data were acquired in data dependent mode with Orbitrap full MS scans collected at 60,000 resolution and the top 15 most intense precursors selected for CID MS/MS fragmentation in the ion trap. MS2 spectra were searched using Mascot, both against a concatenated target-decoy Uniprot database of human proteins as well as against a small database containing wild-type, E545K and H1047R mutant p110α sequences in order to identify mutant peptides. Peptide spectral matches for the Uniprot search were rough filtered using linear discriminant analysis to 10% false discovery rate, then confirmed via manual inspection. Extracted ion chromatograms and peak area integration for p110α peptides were generated with 10 ppm mass tolerances using in-house software (MSPlorer). Peak area data for each of 14 peptides (see Figure 3D and Figures S4C and S4D) were normalized on a per-replicate basis to the most abundant peak area among the six samples. In cases where duplicate injections (technical replicates) were available, normalized data for the two replicates were averaged to generate a single normalized peak area per peptide-condition-experiment for the protein sequence plots (Figure S4C). For statistical analysis, unnormalized peak areas across the four biological replicates were consolidated in R using linear mixed effects modelling (lme4 package) to determine the relative ratio and measures of uncertainty (from which can be derived p-values, confidence intervals, etc) for the comparison of DMSO (vehicle) versus taselisib (500 nM) treatments per cell line for each of total p110α, WT p110α, and mutant p110α (Bates et al., 2015). Total p110α was determined based on the data generated from the following peptides: EATLITIK (residues 39-46; 444.77481 m/z), DLNSPHSR (155-162; 463.22945 m/z), LCVLEYQGK (241-249), VCGCDEYFLEK (254-264; 710.30246 m/z), VPCSNPR (376-382), EAGFSYSHAGLSNR (503-516; 748.35281 m/z), YEQYLDNLLVR (641-651; 713.37540 m/z), FGLLLESYCR (684-693; 629.32042 m/z), and LINLTDILK (712-720; 521.83039 m/z). For cells bearing the H1047R mutation (i.e. HCC-1954), WT p110α was determined based on the QMNDAHHGGWTTK (1042-1054; 749.82824 m/z) peptide and mutant p110α based on QMNDAR (1042-1047; 375.66360 m/z) and HGGWTTK (1048-1054; 393.6983 m/z) peptides. For cells bearing the E545K mutation (i.e. HCC-202), WT p110α was determined based on the DPLSEITEQEK (538-548; 644.81917 m/z) peptide and mutant p110α based on DPLSEITK (538-545; 451.74627 m/z) peptides. Peptides from these two mutant loci were not considered when determining total p110α. Whenever applicable, cysteine residues within p110α peptides were analysed in their carbamidomethylated form (+57.0215 Da) and methionine residues quantified based on their singly oxidized (Met630 sulfoxide) form (+15.9949 Da). Quantified peptide peak areas observed after treatment with taselisib or DMSO were analysed to estimate differential abundances and accompanying measures of uncertainty for total, WT, and mutant, respectively, across the two treatments. These three fractional abundances were combined to estimate the relative proportions (and associated 95% confidence intervals) of WT versus mutant in each mutant-containing cell line in both the DMSO and taselisib treated conditions by applying the conservation of mass principle, as follows. Suppose “p” and “1-p” denote the fractions of WT and mutant p110α, respectively, in a DMSO-treated cell culture. Comparing peptides quantified after treatment with taselisib or DMSO from the total, WT-only, and mutant-only categories detailed above, define the three fractions: f_total = fraction of total p110α remaining after taselisib, relative to DMSO f_WT = fraction of WT p110alpha remaining after taselisib, relative to DMSO f_mut = fraction of mutant p110α remaining after taselisib, relative to DMSO. The conservation of mass principle requires that f_total = f_WT * p + f_mut * (1-p). Solving for “p” (the fraction of WT p110α in DMSO-treated cells) yields an estimate of p = (f_total – f_mutant) / (f_WT - f_mutant) where log-scale estimates of the three fractions {f_total, f_WT, f_mutant} are obtained from the linear mixed effects model. All other quantities of interest (with estimated standard errors, confidence intervals, p-values, etc.) likewise can be estimated as functions of these three fractions and their measures of uncertainty, which are also obtained from the linear mixed effects model. Data are plotted as relative intensity values normalized to 1.0 and error bars represent the 95% confidence intervals for each measurement based on the linear effects model.

## Acknowledgements

Support for third-party editing assistance for this manuscript, furnished by Daniel Clyde, PhD, of Health Interactions, was provided by F. Hoffmann-La Roche Ltd.

---

**Figure S1.**
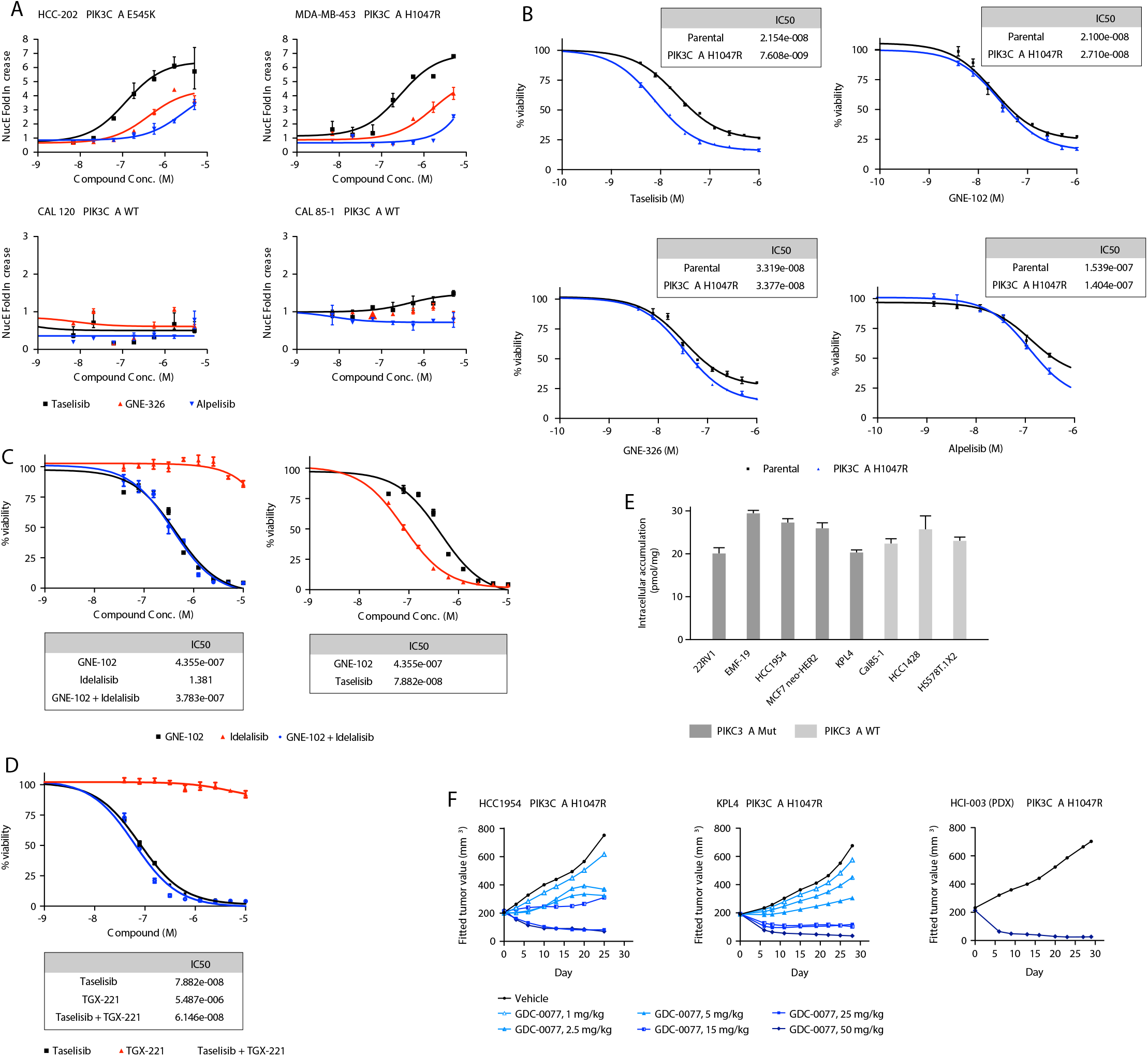
Taselisib and GDC-0077 have increased potency in *PIK3CA*-mutant cancer cells. **(A)** PI3K inhibitors assessed for apoptosis induction in mutant and wild-type breast cancer cell lines in 72 hour Nucleosome ELISA. Error bars are standard deviation of triplicates. **(B)** Cell potency in 4-day CellTiter-Glo**®** viability assay for taselisib and PI3Kα inhibitors BYL719, GNE-102, GNE-326 in SW48 isogenic H1047R and wild-type cells. Error bars are standard deviation of triplicates. **(C)** Combination of PI3Kα inhibitor GNE-102 with PI3Kδ inhibitor idelalisib in 4-day viability assay in HCC1954 PIK3CA H1047R mutant cells. Error bars are standard deviation of quadruplicates. **(D)** Combination of taselisib with PI3Kβ inhibitor TGX-221 in HCC1954 PIK3CA H1047R mutant cells in a 4-day viability assay. Error bars are standard deviation of quadruplicates. **(E)** Intracellular drug concentrations of taselisib in cancer cell lines treated for 18 hours with 1 *µ*M taselisib. Results of LC/MS/MS for triplicate wells are shown. Error bars are standard deviation of triplicates. **(F)** *In vivo* efficacy of GDC-0077 in PIK3CA H1047R breast cancer xenograft HCC1954 and KPL4 models, and breast cancer HCI003 PDX (patient-derived xenograft) model. GDC-0077 dosed orally and daily (QD) in MCT (0.5 % methycellulose/0.2% Tween-80) vehicle.

**Figure S2:**
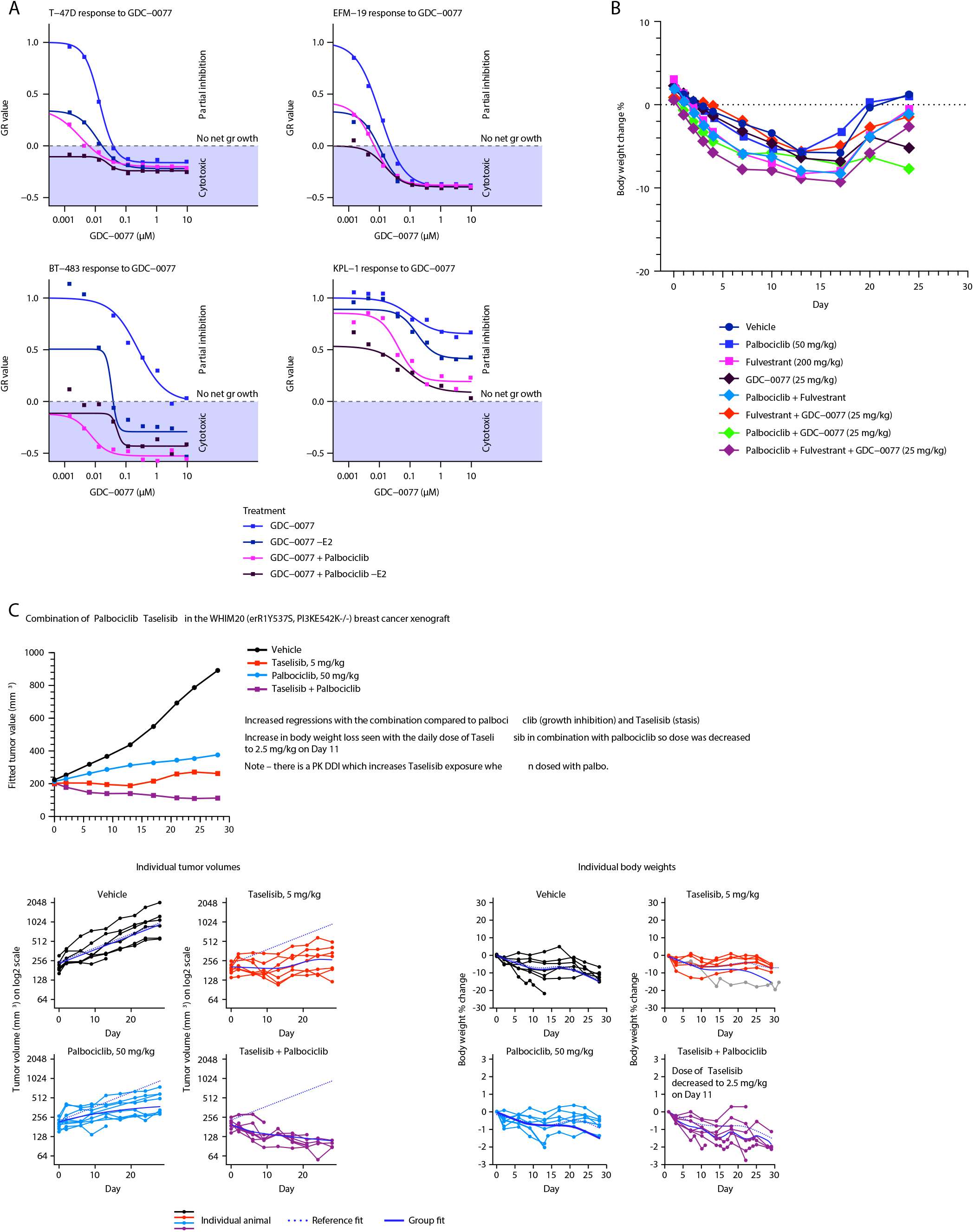
Activity of GDC-0077 in combination with Palbociclib and/ or Fulvestrant. **(A)** Dose response curve of MCF-7 cells treated with GDC-0077 alone (blue), without E2 to mimic aromatase inhibitor (dark blue). **(B)** Weight loss for MCF-7 xenograft experiment. **(C)** Efficacy and weight loss for MDA-MB-453 xenograft experiment.

**Figure S3.**
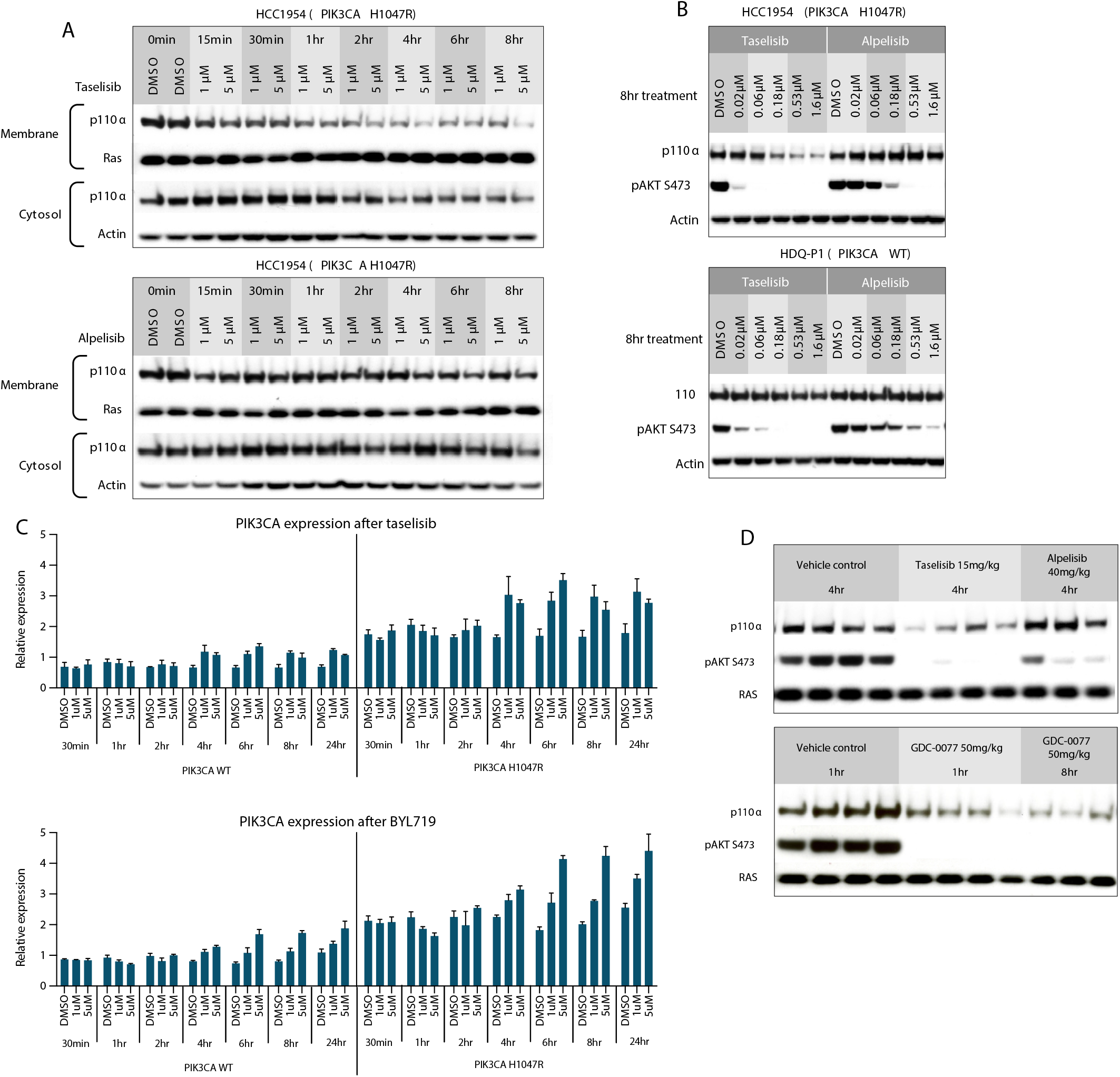
Taselisib and GDC-0077 induce mutant p110α and not WT p110a degradation. **(A)** Subcellular fractionation of HCC1954 PIK3CA H1047R mutant cells treated with PI3K inhibitors taselisib or BYL719 for up to 8 hours, followed by p110α western blot. **(B)** Taselisib and BYL719 treatment for 8 hours in HCC1954 PIK3CA H1047R mutant cells and HDQ-P1 *PIK3CA*-wild-type cells. **(C)** Quantitative RT-PCR shows no reduction in *PIK3CA* RNA expression in HCC1954 cells treated with taselisib or BYL719 for up to 24 hours. Error bars are standard deviation of triplicates. **(D)** Membrane-associated p110a expression following single oral dose of 15 mg/kg taselisib, 40 mg/kg BYL719, or 50 mg/kg GDC-0077 treatment in HCC1954 xenograft tumors.

**Figure S4.**
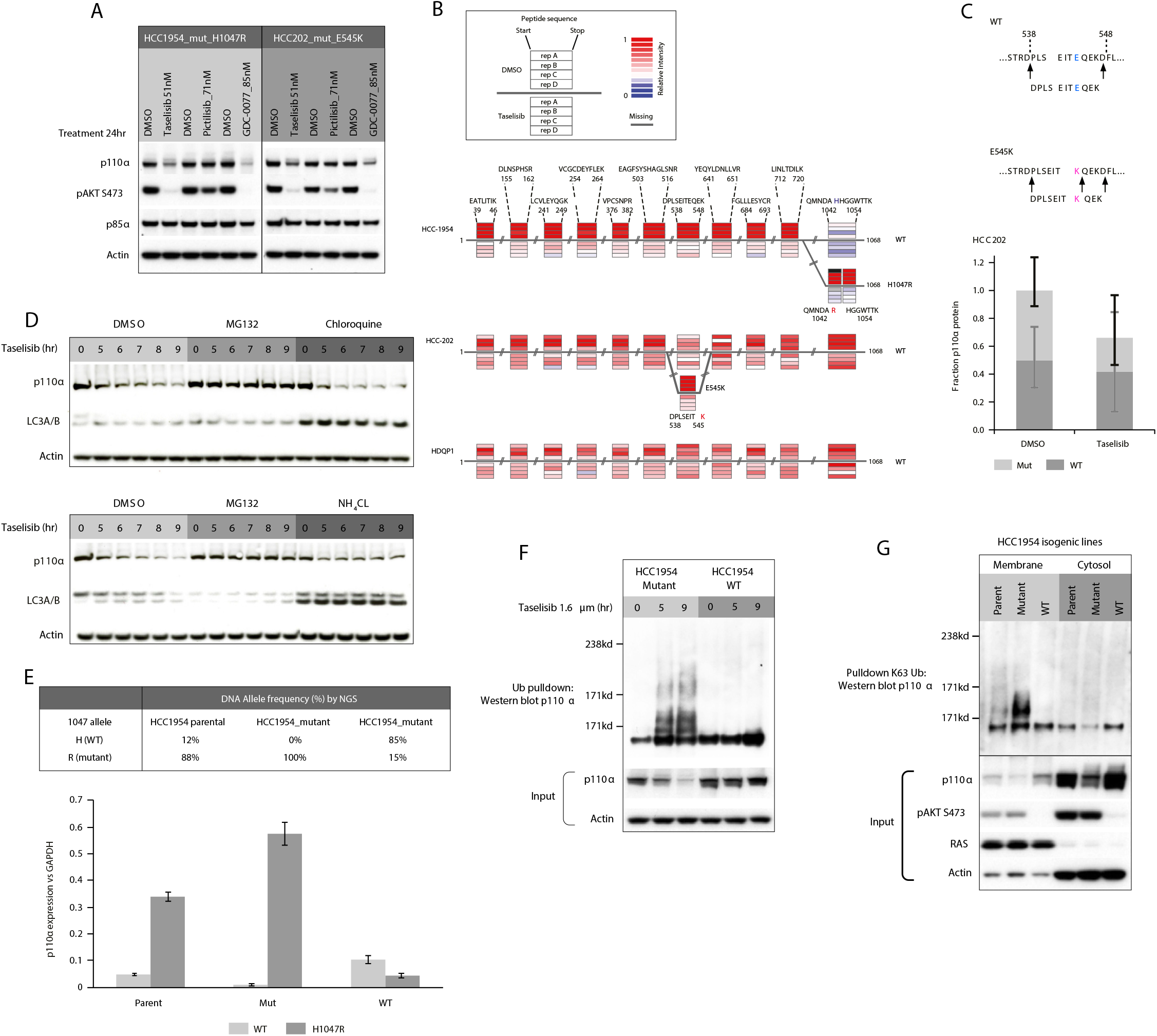
Taselisib depletes mutant p110α protein through ubiquitin and proteasome mechanism in a dose and time dependent manner. **(A)** *PIK3CA*-mutant cells were treated with PI3K inhibitors at concentrations relevant to plasma concentrations achieved with clinically administered doses for pictilisib (GDC-0941) (330 mg), taselisib (6 mg), and GDC-0077 (9 mg). **(B)** Quantitative mass spectrometry analysis of p110α levels in HCC1954, HCC202 and HDQP1 cells treated with either DMSO or 500 nM taselisib for 24 hours (n=4). For each cell line, the relative abundance of peptides from loci spanning the protein are compared between DMSO (above line) and taselisib-treated (below line) cells. For mutant loci (H1047R in HCC1954 cells, E545K in HCC202 cells), the protein sequence is split to depict the wild-type and mutant-specific peptides in parallel. **(C)** Mass spectrometry analysis of peptides representing wild-type and E545K mutant p110α from HCC202 breast cancer cells treated for 24 hours with either DMSO or 500 nM taselisib. The neo-trypic peptide generated from p110α E545K was used to assess mutant protein levels compared to wild-type protein in the same lysate (n=4). **(D)** Lysosomotropic agents chloroquine and ammonium chloride (NH_4_Cl) do not rescue p110α degradation induced by 1.6 *µ*M taselisib in HCC1954 cells **(E)** NGS and qRT-PCR for allele-specific mRNA expression in HCC1954 parental and isogenic HCC1954_mutant and HCC1954_wild-type cells. Error bars are standard deviation of triplicates. **(F)** HCC1954_mutant and HCC1954_wild-type isogenic cells were treated with 1.6 *µ*M taselisib for up to 9 hours. Pull down of ubiquitinated protein was followed by western blotting with anti-p110α antibody. **(G)** Subcellular fractionation of HCC1954_parental and isogenic HCC1954_mutant and HCC1954_WT cells. Pull down of ubiquitinated protein was followed by western blotting with anti-p110alpha.

**Figure S5.**
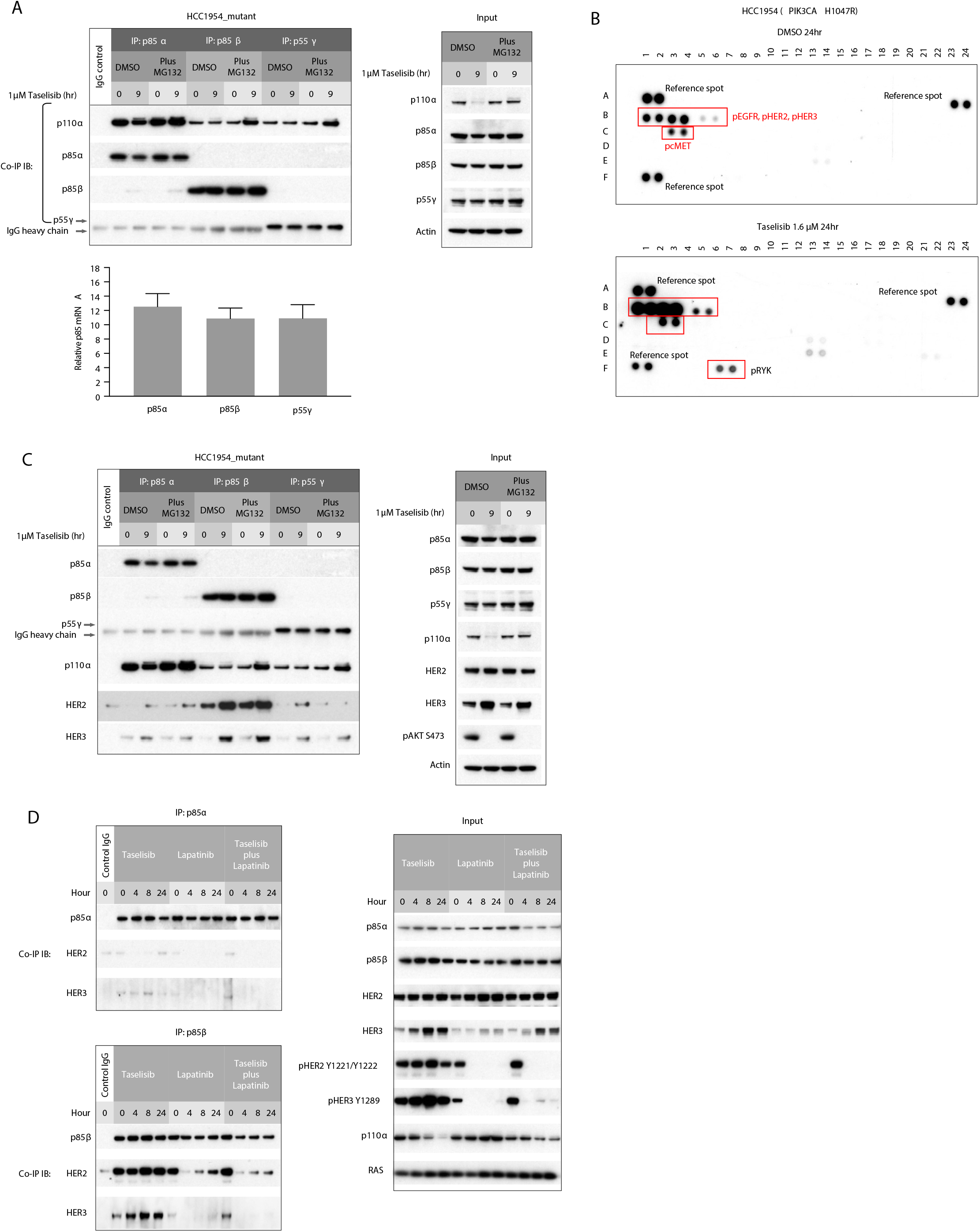
Taselisib-mediated degradation of mutant p110α occurs preferentially at the plasma membrane. **(A)** HCC1954_mutant cells were treated with 1 *µ*M taselisib alone or combination with proteasome inhibitor MG132. Cell lysates were precipitated with p85α, p85β or p55γ antibody, followed by immunoblot with antibodies indicated to the left. Real time qPCR assays of the p85 isoforms in RNA collected from untreated cells. **(B)** HCC1954 cells were treated with 1.6 *µ*M taselisib for 24 hours as indicated. Cell lysates were prepared and total protein were applied to pRTK arrays. Arrows indicate RTKs whose phosphorylation was up-regulated following the treatment. **(C)** HCC1954_mutant cell line was treated with 1 *µ*M taselisib alone or in combination with proteasome inhibitor MG132. Cell lysates were precipitated with p85α, p85β or p55γ antibody, followed by immunoblot with antibodies indicated to the left. **(D)** HCC1954_mutant cell line was treated with 1 *µ*M taselisib alone or in combination with lapatinib. Cell lysates were precipitated with p85α or p85β antibody, followed by immunoblot with antibodies indicated to the left.

**Figure S6.**
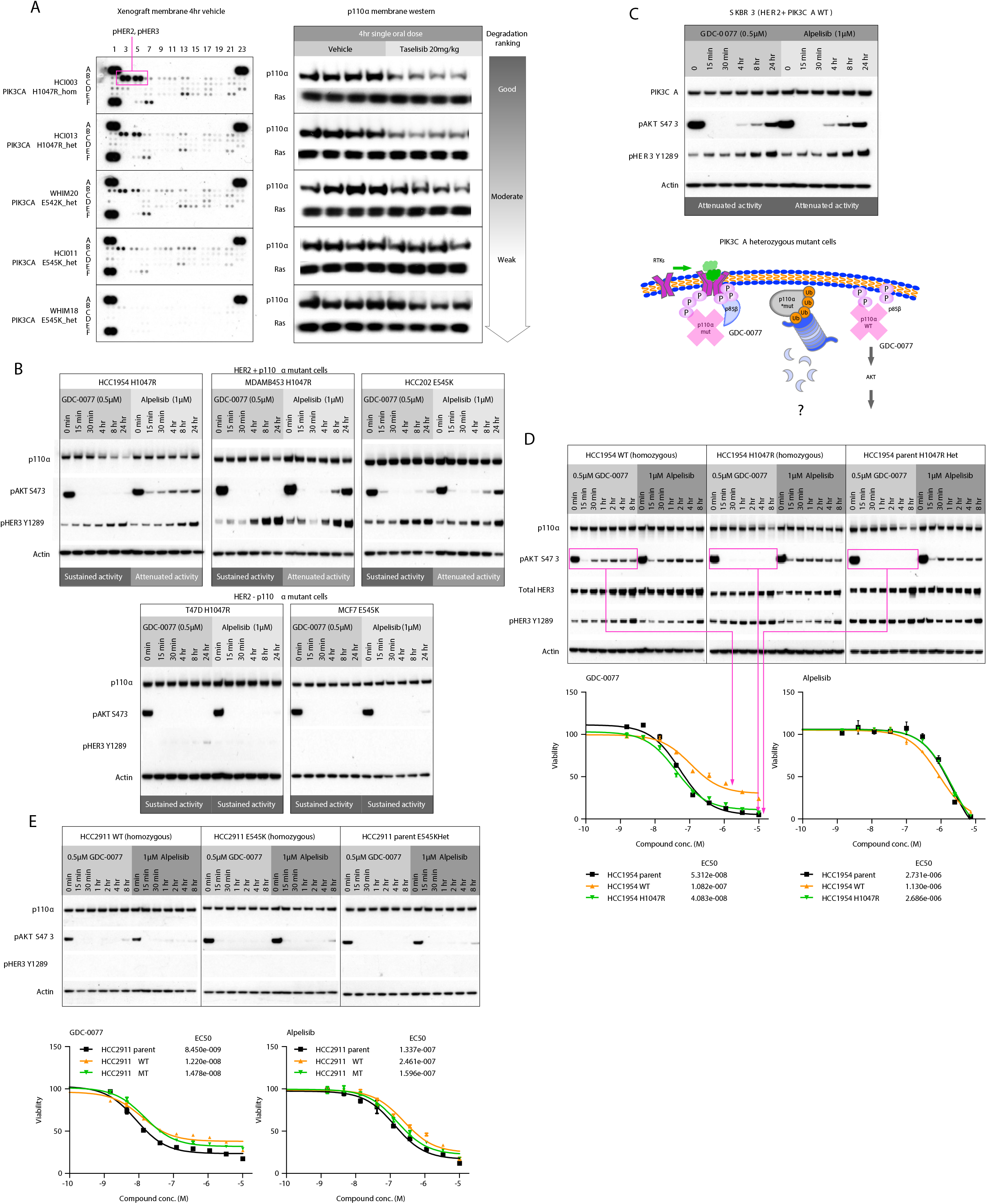
RTK activity plays a key role in regulating p110α degradation. **(A)** Western blot for membrane associated p110α in a subset of treated (taselisib 20 mg/kg 4hr single dose) versus untreated PDX tumors HCI-003, HCI-013, WHIM20, HCI-011, WHIM18. Total lysates from untreated tumor were applied to pRTK arrays. **(B)** Cell lines were harvested after treatment with 0.5 *µ*M GDC-0077 or 1 *µ*M BYL719 at various time points followed by western blot analysis with indicated antibodies **(C)** Mechanistic model of drug-induced p110α degradation in heterozygous *PIK3CA*-mutant cells. SKBR3 (HER2-positive *PIK3CA*-wild-type) cells were treated with 0.5 *µ*M GDC-0077 or 1 *µ*M BYL719 at various time points followed by western blot analysis with indicated antibodies. **(D)** HCC1954 parental and *PIK3CA*-wild-type and -mutant isogenic cell lines are harvested after treatment with 0.5 *µ*M GDC-0077 or 1 *µ*M BYL719 at various time points followed by western blot analysis with indicated antibodies. Cell viability IC_50_ values determined by quantifying ATP from same cell lines at 5 days post-treatment. **(E)** HCC2911 parental and *PIK3CA*-wild-type and -mutant isogenic cell lines are harvested after treatment with 0.5 *µ*M GDC-0077 or 1 *µ*M BYL719 at various time points followed by western blot analysis with indicated antibodies. Cell viability IC_50_ values determined by quantifying ATP from same cell lines at 5 days posttreatment.

